# OGT controls mammalian cell viability by regulating the proteasome/mTOR/mitochondrial axis

**DOI:** 10.1101/2021.09.19.461012

**Authors:** Xiang Li, Xiaojing Yue, Hugo Sepulveda, Rajan A. Burt, David A. Scott, Steven A. Carr, Samuel A. Myers, Anjana Rao

## Abstract

*O*-GlcNAc transferase (OGT) catalyzes the modification of serine and threonine residues on nuclear and cytosolic proteins with *O*-linked N-acetylglucosamine (GlcNAc). OGT is essential for mammalian cell viability, but the underlying mechanisms are still enigmatic. We employed a genome-wide CRISPR-Cas9 viability screen in mouse embryonic stem cells (mESCs) with inducible *Ogt* gene deletion and showed that the block in cell viability induced by OGT deficiency stems from mitochondrial dysfunction secondary to mTOR hyperactivation. In normal cells, OGT maintains low mTOR activity and mitochondrial fitness through suppression of proteasome activity; in the absence of OGT, increased proteasome activity results in increased steady-state amino acid levels, which in turn promote mTOR lysosomal translocation and activation, and increased oxidative phosphorylation. mTOR activation in OGT-deficient mESCs was confirmed by an independent phosphoproteomic screen. Our study highlights a novel series of events whereby OGT regulates the proteasome/ mTOR/ mitochondrial axis in a manner that maintains homeostasis of intracellular amino acid levels, mitochondrial fitness and cell viability. A similar mechanism operates in CD8^+^ T cells, indicating its generality across mammalian cell types. Manipulating OGT activity may have therapeutic potential in diseases in which this signaling pathway is impaired.

## Introduction

*O*-GlcNAc transferase (OGT) is an essential X-chromosome-encoded enzyme that catalyzes the addition of *N*-acetylglucosamine (GlcNAc) to the hydroxyl groups of serine and threonine residues on many nuclear and cytosolic proteins^1,2^. This posttranslational modification is reversible and is actively removed by the *O*-GlcNAcase OGA^1,2^. It has been known for more than two decades that OGT is essential for mammalian cell viability^3^, but the underlying mechanisms are still unclear^1^. Given the close association between OGT and human diseases such as cancer, diabetes and cardiovascular disease^4-6^, knowledge of the mechanisms by which OGT controls cell viability is essential to understand the role of OGT and the *O*-GlcNAc modification in cellular function.

A functional interaction of OGT with mitochondria has been suggested by several previous studies, many of which focused on mitochondrial OGT (mOGT), a splice variant of OGT that contains a mitochondrial targeting sequence^7^. Detailed analyses of mOGT function have been complicated by difficulties in specifically and efficiently depleting mOGT^8^ and the reportedly low levels of *O*-GlcNAcylated proteins in the mitochondria^9^. The relation between *O*-GlcNAcylation and mitochondrial function depends strongly on the particular cellular system used. Increasing O-GlcNAc levels by treatment with the OGA inhibitor thiamet-G led to increased oxygen consumption and ATP production rates in cardiac mitochondria^10^, but decreased basal oxygen consumption rates (OCR) and ATP production without an effect on maximal OCR in SH-SY5Y neuroblastoma or NT2 human embryonal carcinoma cells^11^. Partial (50-70%) depletion of total OGT with a pan-OGT siRNA, which would be expected to decrease cellular *O*-GlcNAc levels, resulted in increased basal and maximal oxygen consumption rates (OCR) in HeLa cells^8^, whereas hematopoietic stem cells (HSCs) in which the *Ogt* gene was deleted by treatment of *Ogt floxed Mx1Cre* mice with polyI:polyC had increased mitochondrial mass coupled to decreased copy numbers of mitochondrial DNA and decreased spare respiratory capacity, defined as the difference between maximal and basal OCR^12^. However, these findings are complicated by the fact that *Ogt* deletion resulted in severe reduction of the numbers of HSC, and likely promoted the accumulation of cells in which *Ogt* deletion was incomplete. The effects of OGT deletion on mitochondrial function need to be reevaluated in better-defined cellular systems with efficient OGT deletion.

The mechanistic target of rapamycin (mTOR) is a serine/threonine kinase that regulates fundamental cellular processes including cell growth and metabolism in response to environmental and intracellular signals^13,14^. mTOR is the catalytic subunit of two functionally distinct protein complexes, mTOR complex 1 (mTORC1) defined by the presence of the rapamycin-sensitive subunit Raptor, and mTORC2, which contains the alternate subunit Rictor^15^. The mTORC1 complex has been described as a “coincidence detector” because it is activated only when both growth factors and nutrients are present to facilitate cellular growth^13^. mTORC1 activation requires two separate steps that occur after translocation of mTOR to the lysosomal membrane^13,14^. Growth factors and cellular stress signals induce activation of the small GTPase Rheb, which is located on the lysosomal membrane and directly stimulates mTOR kinase activity in its GTP-bound form. However, mTORC1 only localizes in the vicinity of GTP-bound Rheb when nutrient signals such as amino acids are available to activate the Rag complex, a heterodimer of the small GTPases, RagA and RagC (or RagB and RagD). When RagA (or RagB) are GTP-bound and RagC (or RagD) are bound to GDP, the Rag heterodimers recruit mTORC1 to the lysosomal membrane. The Rag heterodimer is itself activated and anchored to the lysosomal membrane by the pentameric Ragulator complex, a non-canonical guanine nucleotide exchange factor that contains Lamtor1-5^16,17^. Thus, mTOR indirectly senses amino acid levels through proteins associated with the Rag and Ragulator complexes. Moreover, mTORC1 has been reported to regulate mitochondrial function in at least two ways: through transcriptional control of mitochondrial regulators such as PGC-1α (PPARγ coactivator 1α)^18^, and by regulation of protein synthesis of a subset of mitochondrial proteins encoded by nuclear DNA through inhibition of eukaryotic translation initiation factor 4E-binding proteins (4E-BPs)^19^.

To investigate why OGT is essential for the survival of proliferating cells^1^, we generated mouse embryonic stem cell (mESC) lines with one or two floxed *Ogt* alleles respectively. The blastocysts from which the ESCs were derived also expressed a Cre-ERT2 fusion protein for inducible deletion of the *Ogt* gene, and a *Rosa26-YFP*^*LSL*^ reporter gene to mark cells in which Cre-ERT2 had been activated in the nucleus with 4-hydroxytamoxifen (4-OHT). We used these cells to perform genome-wide CRISPR-Cas9 screens for small guide RNAs (sgRNAs) that rescued the arrested cell proliferation of *Ogt*-deleted mESCs. The enriched sgRNAs targeted a large number of genes related to mitochondrial function, as well as genes whose products were involved in amino acid sensing by mTOR. We show that OGT deficiency leads to arrested cell proliferation and loss of mESC viability at least partly by increasing proteasome activity and intracellular amino acid levels, which in turn promotes striking lysosomal translocation and activation of mTORC1. The net result is a deleterious increase in mitochondrial oxidative phosphorylation (OXPHOS), which correlates strongly with decreased cell viability. Genome-wide proteomic and phosphoproteomic analyses show extensive changes in global signalling and confirm our finding of mTOR hyperactivation in OGT-deficient cells. By highlighting the biochemical connection between OGT deficiency and high aberrant activation of proteasome function, the mTOR pathway, and mitochondrial OXPHOS, our studies provide important mechanistic insights into the role of *O*-GlcNAc modifications in intracellular homeostasis and cell viability.

## Results

### Inducible deletion of the *Ogt* gene impairs cell viability in mESCs

To overcome the lethality of *Ogt* deletion in mouse embryonic stem cells (mESCs)^3^, we generated conditional *Ogt floxed* mice homozygous for knock-in alleles of *Cre-ERT2* or *LSL-YFP* in the *Rosa26* locus. We crossed *Ogt*^*floxed*^*Cre-ERT2*^*KI/KI*^ mice with *Ogt*^*floxed*^*Rosa26-LSL-YFP*^*KI/KI*^ mice to obtain *Ogt*^*floxed*^*Cre-ERT2*^*+/KI*^*Rosa26-LSL-YFP*^*+/KI*^ (hereafter termed *Ogt fl*) blastocysts (embryonic day 3.5), from which we derived *Ogt fl* mESC lines (**Fig. 1a**). Subsequent experiments were performed with male *Ogt fl* mESCs to maximize the likelihood of complete *Ogt* gene deletion in response to 4-hydroxytamoxifen (4-OHT) treatment.

**Fig. 1.**
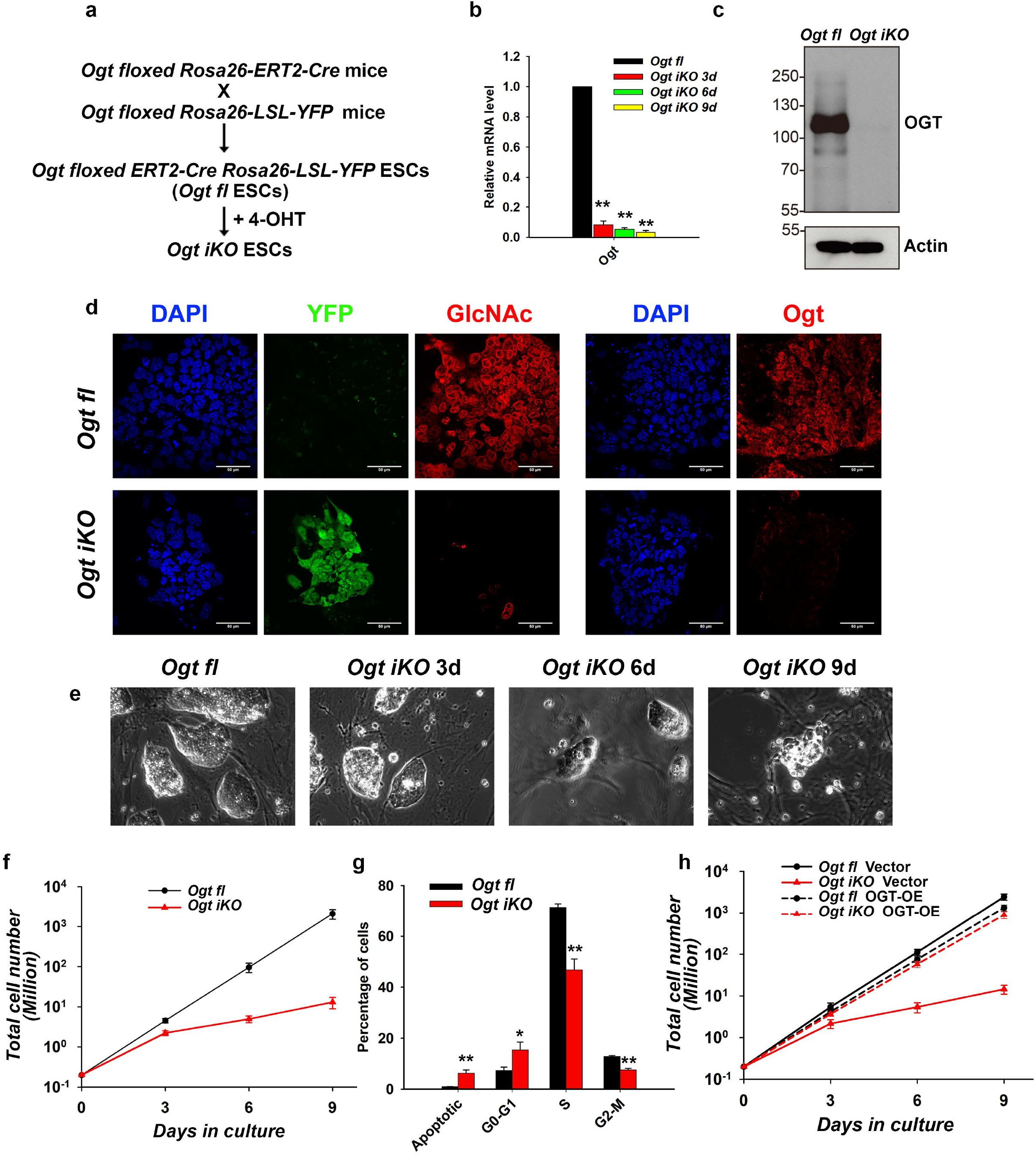
Generation of mESCs with inducible *Ogt* deletion. **a**, Scheme used for the generation of inducible *Ogt fl Cre-ERT2+/KI Rosa26-YFP+/KI* (*Ogt* fl) mESC lines. **b**, Quantitative real-time PCR (qRT-PCR) analysis of *Ogt* transcript levels in *Ogt fl* control and *Ogt iKO* mESCs 3, 6, 9 days after 4-OHT treatment. The expression level of *Ogt* mRNA is shown relative to the level in control mESCs. Data are shown as mean ± SD (N=3). **c**, Western blot analysis of OGT protein level in *Ogt fl* mESCs treated with or without 4-OHT for 6 days. Actin was used as a loading control. **d**, Immunohistochemistry of *Ogt fl* and *Ogt iKO* mESCs 6 days after 4-OHT treatment, using antibodies against *O*-GlcNAc and OGT. Nucleus staining: DAPI (blue). Scale bar: 10 μm. **e**, Phase contrast images of *Ogt fl* control and *Ogt iKO* mESCs 3, 6, 9 days after 4-OHT treatment. **f**, Cumulative growth curves of *Ogt fl* mESCs treated with or without 4-OHT at day 0 and counted at each passage (every 3 days) thereafter until day 9. Data are shown as mean ± SD (N=3). **g**, Percentage of *Ogt fl* and *Ogt iKO* mESCs 6 days after 4-OHT treatment in different phases of the cell cycle. Data are shown as mean ± SD (N=3). Note the logarithmic scale on the Y-axis. **h**, Cumulative growth curves of *Ogt fl* mESCs stably expressing empty vector or wildtype OGT treated with or without 4-OHT. Data are shown as mean ± SD (N=3).

At day 3 after 4-OHT treatment, the extent of *Ogt* gene deletion was >90% as assessed by qRT-PCR for *Ogt* mRNA, and the extent of deletion continued to increase at day 6 and 9 after 4-OHT treatment (**Fig. 1b**). Deletion at the protein level was nearly complete by 6 days as assessed by western blot (**Fig. 1c**). Immunostaining confirmed that both OGT and its catalytic product, the *O*-GlcNAc modification, were almost completely eliminated in *Ogt*-deleted cells (hereafter termed *Ogt iKO* mESCs; i for “inducible”) by day 6 (**Fig. 1d**). *Ogt iKO* mESCs gradually underwent a significant morphology change, from large, tightly packed colonies in the case of *Ogt fl* mESCs to small, loosely distributed colonies after *Ogt* deletion (**Fig. 1e**). OGT-deficient mESCs displayed a significantly impaired growth rate beginning at ∼3 days after 4-OHT addition, resulting in an ∼90% decrease in the number of *Ogt iKO* cells by day 6, and >95% decrease by day 9 (**Fig. 1f**, please note logarithmic scale of Y-axis). The *Ogt iKO* mESCs also showed decreased cell viability, and decreased expression of the pluripotency markers *Oct4* and *Nanog* (**Extended Data Fig. 1a, b**). Cell cycle analysis using BrdU, a thymidine analogue that is incorporated into newly synthesized DNA during S phase, and 7-AAD, a DNA intercalating dye, showed that *Ogt iKO* mESCs displayed significant cell cycle arrest at the G0-G1 stage as well as a significant increase of apoptotic cells by day 6 (**Fig. 1g, Extended Data Fig.1c**). G1-phase arrest of cell cycle progression provides an opportunity for cells to repair DNA and other damage or if this fails, to undergo apoptosis^20^. The impaired growth rate was due specifically to *Ogt* deletion, since it was rescued by overexpression of wildtype (WT) OGT in *Ogt iKO* mESCs, concomitantly with restoration of *O*-GlcNAc staining (**Fig. 1h, Extended Data Fig. 1d**). 4-OHT treatment did not affect the cell growth of WT mESCs (**Extended Data Fig. 1e**).

### A genome-wide CRISPR-Cas9 screen identifies key regulators of cell survival in OGT-deficient mESCs

To identify genes whose deletion could restore cell proliferation and increase the survival of *Ogt iKO* mESCs, we conducted an unbiased genome-wide CRISPR-Cas9 viability screen. We stably expressed Cas9 in *Ogt fl* mESCs, and lentivirally transduced them with the sgRNA Brie library^21^, which contains a pool of 78,637 sgRNAs targeting 19,647 genes with 4 sgRNAs per gene and 1000 control non-targeting sgRNAs. After puromycin treatment for 7 days to select for cells stably expressing the sgRNAs, the cells were split into two groups, an untreated (*Ogt fl*) group that served as the control, and a second group treated with 4-OHT for 6 days to induce *Ogt* deletion (*Ogt iKO*). On day 13, YFP-negative cells from the control *Ogt fl* group and YFP^+^ cells from the *Ogt iKO* group were sorted, and the enriched sgRNAs were identified by next generation deep sequencing (**Fig. 2a**). The sequencing data were analyzed using the PinAPL-Py platform^22^.

**Fig. 2.**
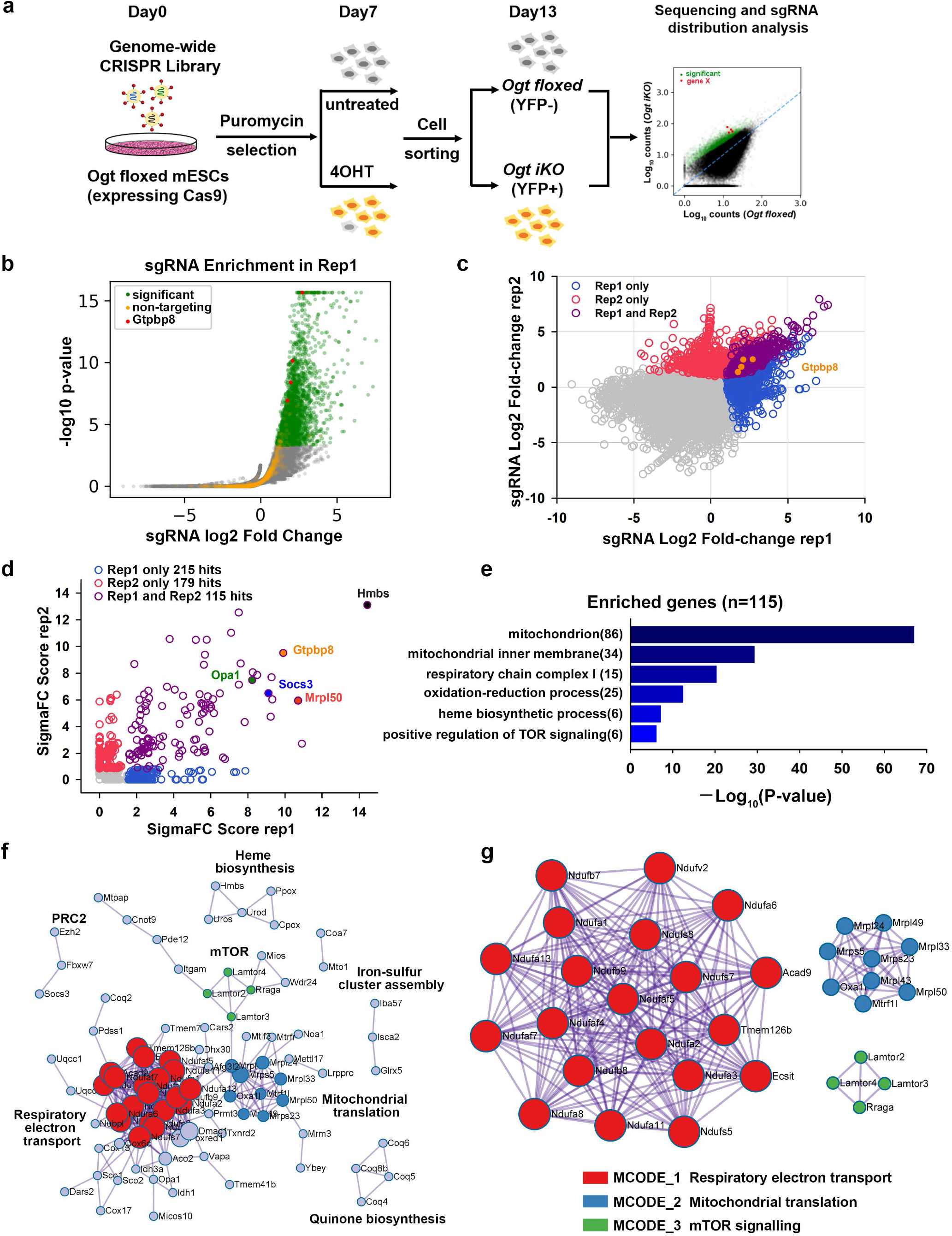
Genome-wide CRISPR-Cas9 screen to identify key regulators of cell survival in OGT-deficient mESCs. **a**, Strategy for the genome-wide CRISPR-Cas9 viability screen to identify key regulators of cell growth in OGT-deficient mESCs. **b**, Volcano plot of p-value versus fold enrichment for each sgRNA in Replicate 1. Significantly enriched sgRNAs are shown in green and non-targeting control sgRNAs in orange. All four sgRNAs for *Gtpbp8* were significantly enriched in the screen (red dots). **c**, sgRNAs that were enriched in one or both biological replicates of the CRISPR-Cas9 viability screen. The four sgRNAs targeting *Gtpbp8* are highlighted. The correlation between the fold changes in sgRNA abundance has a Pearson’s r of 0.438. **d**, Rank of enriched hits by gene score, calculated based on the log fold changes of all sgRNAs targeting each gene and the number of sgRNAs targeting each gene that reached statistical significance. 115 enriched hits in both biological replicates are shown in purple. A few of the most highly scored hits (*Hmbs, Gtpbp8, Opa1, Socs3* and *Mrpl50*) are labeled. **e**, Gene ontology analysis of biological pathways of the 115 enriched hits. **f**, Metascape visualization of the protein-protein interactome network of the 115 highly-scored hits from the CRISPR screen. Each MCODE complex identified by Metascape was assigned a unique color. **g**, Three MCODE complexes and their associated functional pathways identified by Metascape.

We performed two independent screens that were sequenced to a depth of >10M mapped reads per sample; >98% of all genes in the sgRNA library were represented in each screen (**Extended Data Fig. 2a**). The volcano plot showed significantly enriched sgRNAs in green and 1000 control non-targeting sgRNAs in orange which allowed us to calculate an empirical false discovery rate of 3.1% for replicate1 and 1.1% for replicate 2 (**Fig. 2b and Extended Data Fig.2b; Supplementary Table 1**). The overlap of enriched sgRNAs and their rank in each replicate is shown in **Fig. 2c**. Gene scores were calculated considering the log fold changes of all sgRNAs targeting each gene and the number of sgRNAs targeting each gene that reached statistical significance^22^. In the two replicate experiments, 115 hits were identified by the overlap of sgRNAs enriched in the *Ogt iKO* groups compared to the control *Ogt fl* groups (p-value < 0.01) (**Extended Data Fig. 2c; Supplementary Table 2**). Hits with the highest scores, including *Hmbs, Mrpl50, Gtpbp8, Socs3* and *Opa1* are highlighted (**Fig. 2d**).

Gene scores were calculated using SigmaFC, which considers the sum of the log fold changes of all sgRNAs targeting each gene, and multiplies this number by the number of sgRNAs that reached statistically significant enrichment^22^. We plotted SigmaFC scores from two independent screen replicates and only investigated those that were significant across both screens. In the two replicate experiments, 115 candidate genes were identified by the overlap of sgRNAs enriched in the *Ogt iKO* groups compared to the control *Ogt fl* groups (p-value < 0.01) (**Extended Data Fig. 2c; Supplementary Table 2**). Genes with the highest scores, including *Hmbs, Mrpl50, Gtpbp8, Socs3* and *Opa1*, are highlighted in **Fig. 2d**.

More than 80 of the 115 gene candidates were annotated by gene ontology analysis as related to the mitochondrion or the mitochondrial respiratory chain complex (**Fig. 2e**). Metascape, which applies the MCODE (Molecular Complex Detection) algorithm to identify protein complexes^23^, identified three MCODE complexes: respiratory electron transport, including NADH:ubiquinone oxidoreductase family (Nduf) proteins in mitochondrial complex I (*shown in red*); proteins involved in mitochondrial translation, such as mitochondrial ribosomal proteins (Mrpl) (*shown in blue*), and mTOR signaling, specifically Rraga and Lamtor2-4 (*shown in green*) (**Fig. 2f, g**). Metascape analysis also identified protein complexes involved in heme biosynthesis (Hmbs, Uros and Urod), quinone biosynthesis (Coq4, Coq5, Coq6 and Coq8b) and iron-sulfur cluster assembly (Glrx5, Isca2 and Iba57) (**Fig. 2f**). Screen hits related to mitochondria and mTOR signaling are considered in detail below. Other candidate hits are being investigated further as part of a separate project.

### OGT deficiency results in mitochondrial dysfunction

To follow up on the large number of enriched sgRNAs targeting mitochondrial proteins, we assessed mitochondrial function in control and *Ogt*-deleted cells by evaluating mitochondrial membrane potential, mitochondrial mass and mitochondrial oxidative phosphorylation (OXPHOS). Immunostaining and flow cytometry analysis showed that mitochondrial membrane potential assessed by MitoTracker red CMXRos staining (**Fig. 3a-c**), and mitochondrial mass assessed by MitoTracker deep red staining (**Extended Data Fig. 3a-c**), were both increased in *Ogt iKO* relative to *Ogt fl* mESCs. However, OGT deficiency did not affect mitophagy, a process for removal of damaged mitochondria through autophagy, as assessed by using a small-molecule fluorescent probe^24^ (Mtphagy Dye; **Extended Data Fig. 3d-e**). Evaluation of mitochondrial OXPHOS showed that basal and maximal oxygen consumption rates (OCR) as well as ATP production, the indicator of OXPHOS, were aberrantly increased in *Ogt iKO* compared to *Ogt fl* mESCs (**Fig. 3d, Extended Data Fig. 3f)**. As expected, overexpression of wildtype (WT) OGT in *Ogt iKO* mESCs rescued the aberrant increase of mitochondrial OXPHOS (**Extended Data Fig. 3g, h**). OGT deficiency was associated with an approximately 2-fold increase in the levels of reactive oxygen species (ROS) measured by cellROX deep red, presumably as a consequence of increased OXPHOS (**Extended Data Fig. 3i, j)**. However, the ROS scavengers N-acetylcysteine and Vitamin C failed to rescue the viability of OGT-deficient mESCs (data not shown), possibly because mitochondrial OXPHOS but not ROS was required for the activation of apoptotic marker gene Bax and cell death^25^.

**Fig. 3.**
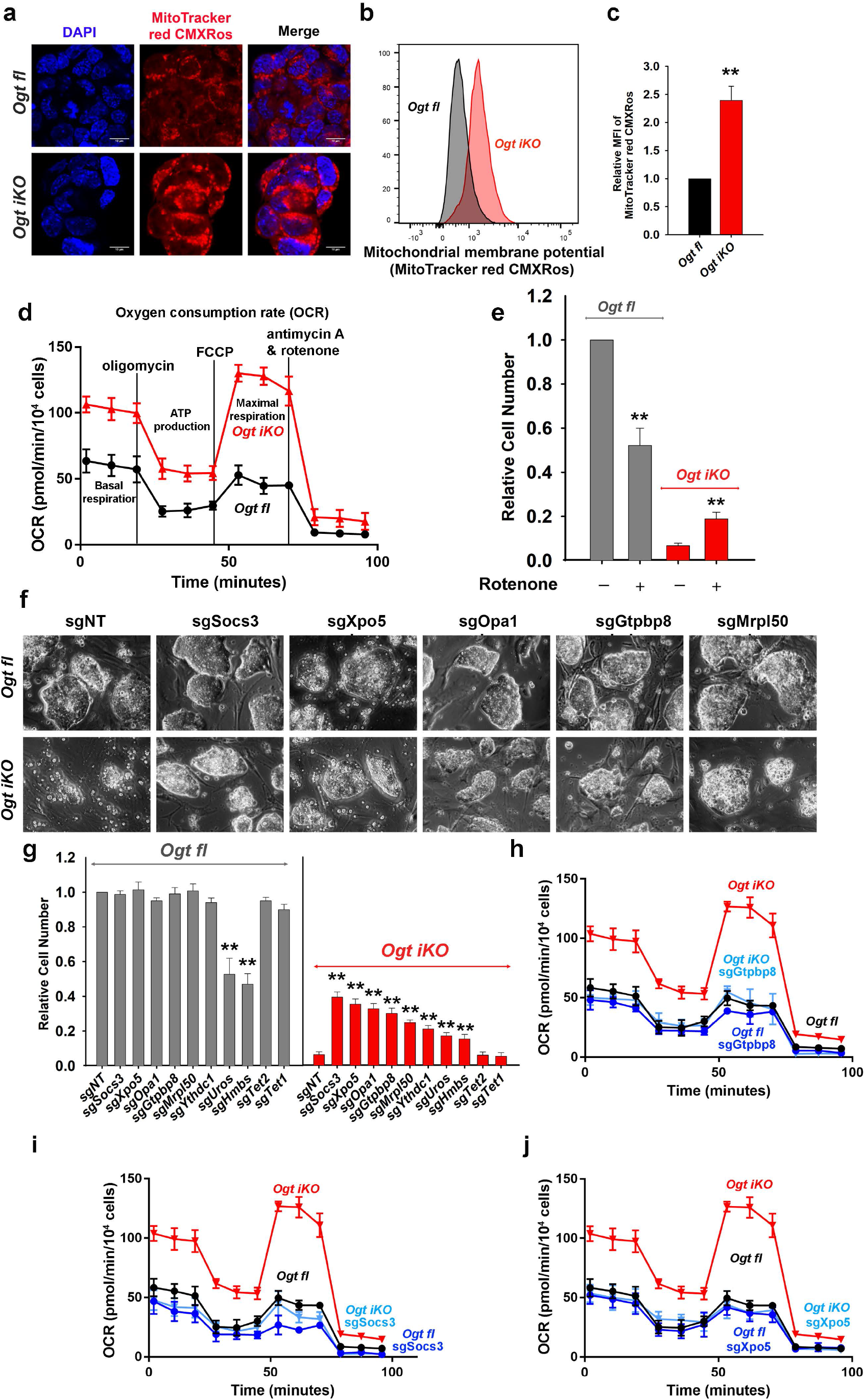
OGT deficiency results in mitochondrial dysfunction. **a**, Fluorescence images of *Ogt fl* and *Ogt iKO* mESCs 6 days after 4-OHT treatment. Cells were stained with MitoTracker red CMXRos. Nucleus staining: DAPI (blue). Scale bar: 10 μm. **b**, Representative histogram of MitoTracker red CMXRos fluorescence in *Ogt fl* and *Ogt iKO* mESCs 6 days after 4-OHT treatment. **c**, Relative MFI (mean fluorescent intensity) of MitoTracker red CMXRos shown in Fig. **3b. d**, Analysis of oxygen consumption rate (OCR) using Seahorse XFe24 in *Ogt fl* and *Ogt iKO* mESCs treated without or with 4-OHT respectively for 6 days. Data are shown as mean ± SD (N=3). **e**, Relative cell numbers of *Ogt fl* and *Ogt iKO* mESCs treated without or with 4-OHT respectively and with or without the mitochondrial Complex I inhibitor rotenone (75 nM) for 8 days. Data are shown as mean ± SD (N=3). **f**, Phase contrast images of *Ogt fl* and *Ogt iKO* mESCs expressing non-targeting sgRNA (sgNT) or sgRNAs targeting the indicated genes (*Socs3, Xpo5, Opa1* and *Gtpbp8*) and treated without or with 4-OHT respectively for 8 days. **g**, Relative cell numbers of *Ogt fl* and *Ogt iKO* mESCs expressing non-targeting sgRNA (sgNT) or sgRNAs targeting the indicated genes (*Socs3, Xpo5, Opa1, Gtpbp8, Mrpl50, Ythdc1, Uros, Hmbs, Tet2* or *Tet1*) and treated without or with 4-OHT respectively for 8 days. Data are shown as mean ± SD (N=3). **h-j**, Analysis of oxygen consumption rate (OCR) using Seahorse XFe24 in *Ogt fl* and *Ogt iKO* mESCs expressing sgRNAs targeting *Gtpbp8* (**h**), *Socs3* (**i**) and *Xpo5* (**j**) and treated without or with 4-OHT respectively for 6 days. Data are shown as mean ± SD (N=3).

Metabolome analyses showed that the TCA cycle metabolites citrate, alpha-ketoglutarate, succinate, fumarate and malate, which are tightly associated with OXPHOS^26^, were increased, while the glycolytic pathway metabolites 3-phosphoglycerate and phosphoenolpyruvate were decreased, in *Ogt iKO* compared to *Ogt fl* mESCs (**Extended Data Fig. 3k**). At the transcriptional level, the majority of genes encoded in the mitochondrial genome showed increased expression levels in *Ogt iKO* mESCs (**Extended Data Fig. 3l**). The mitochondrial complex I inhibitor Rotenone partially rescued the blocked proliferation of *Ogt iKO* mESCs, despite its substantial toxicity for *Ogt fl* mESCs (**Fig. 3e and Extended Data Fig. 4a**), supporting the conclusion that mitochondrial function is hyperactivated in OGT-deficient mESCs. As shown above, reconstitution with WT OGT rescued both the cell proliferation defect and the aberrant increase in mitochondrial OXPHOS of *Ogt iKO* mESCs (**Fig. 1h, Extended Data Fig. 3g, h**), and OGT-reconstituted *Ogt iKO* cells were as sensitive to rotenone as WT cells (50% decrease in cell numbers, compare **Fig. 3e and Extended Data Fig. 4b**).

**Fig. 4.**
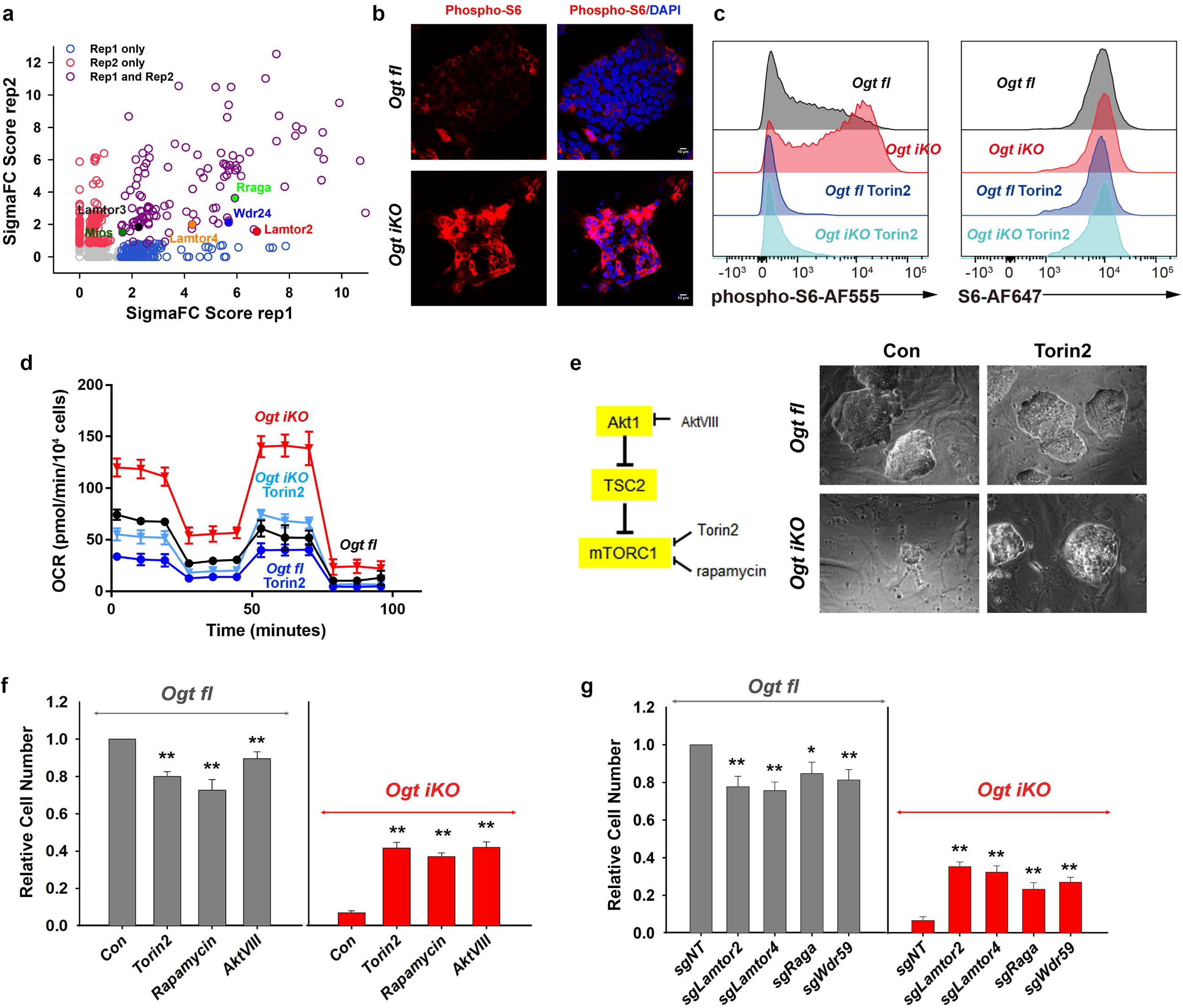
mTOR is hyperactivated in OGT-deficient mESCs. **a**, Scatterplot of enriched genes by gene score. Genes related to positive regulation of mTOR signaling are highlighted. **b**, Immunohistochemistry of *Ogt fl* and *Ogt iKO* mESCs treated without or with 4-OHT respectively for 6 days. Cells were stained with antibody against phospho-S6 ribosomal protein. Nucleus staining: DAPI (blue). Scale bar: 10 μm. **c**, Flow cytometry analysis of phospho-S6 (*left panel*) and total S6 (*right panel*) ribosomal protein in *Ogt fl* and *Ogt iKO* mESCs treated without or with 4-OHT respectively for 6 days and then with or without 200 nM Torin2 for 2 hours. Data are representative of three biological replicates. **d**, Analysis of oxygen consumption rate (OCR) using Seahorse XFe24 in *Ogt fl* and *Ogt iKO* mESCs treated without or with 4-OHT respectively and with or without Torin2 (25 nM) for 6 days. Data are shown as mean ± SD (N=3). **e**, *Left*, Simplified schematic diagram of mTOR pathway activation and the proteins targeted by the inhibitors Torin2 (25 nM), rapamycin (50 nM) and AktVIII (7.5 μM). *Right*, Phase contrast images of *Ogt fl* and *Ogt iKO* mESCs treated with or without Torin2. **f**, Relative cell numbers of *Ogt fl* and *Ogt iKO* mESCs treated without or with 4-OHT respectively and with or without Torin2 (25 nM), Rapamycin (50 nM) and AktVIII (7.5 μM) for 8 days. Data are shown as mean ± SD (N=3). **g**, Relative cell numbers of *Ogt fl* and *Ogt iKO* mESCs expressing gRNAs against the indicated genes (*Lamtor2, Lamtor4, Raga* and *Wdr59*) and treated without or with 4-OHT respectively for 8 days. Data are shown as mean ± SD (N=3).

We used an all-in-one CRISPR/Cas9 vector system to individually interrogate the effects of enriched sgRNAs targeting eight of the top candidate genes identified in our screen (**Fig. 2d**). The candidate proteins that we selected were involve in diverse cellular pathways – mitochondrial proteins (*Opa1, Gtpbp8, Mrpl50*), proteins involved in heme biosynthesis (*Hmbs, Uros*) and proteins involved in other cellular functions (*Socs3, Xpo5, Ythdc1*). Non-targeting (NT) sgRNAs and sgRNAs targeting *Tet1* or *Tet2* (which are expressed in ES cells, but whose sgRNAs were not enriched in our screen) were used as controls. Each gene was efficiently deleted as assessed by qRT-PCR (**Extended Data Fig. 5a**). Transduction with *Cas9* and sgRNAs targeting all 8 genes improved the morphology and growth rate of *Ogt iKO* mESCs, leading to the formation of large, tightly packed colonies as well as promoting a 3- to 7-fold increase in cell numbers; transduction with control NT and *Tet1/Tet2* sgRNAs had no effect (**Fig. 3f, g; Extended Data Fig. 5b**). Moreover, sgRNAs against *Socs3, Xpo5* and *Gtpbp8* restored expression levels of the pluripotency markers *Oct4* and *Nanog* (**Extended Data Fig. 5c**). *Hmbs* and *Uros* sgRNAs reproducibly rescued the proliferation of *Ogt iKO* mESCs to the same degree as rotenone (*compare* **Figs. 3e, 3g**), but were not further considered because they also diminished the proliferation of normal (*Ogt fl*) mESCs (**Fig. 3g**). Notably, sgRNAs targeting the top three candidate genes, *Socs3, Xpo5* and *Gtpbp8*, reduced the aberrant increase of oxygen consumption rates in *Ogt iKO* mESCs to normal levels (**Fig. 3h-j, Extended Data Fig. 6a-c**). Our data indicate that sgRNA-mediated depletion of each of these proteins indirectly inhibits the increase in mitochondrial OXPHOS resulting from OGT depletion, although the exact pathways and mechanisms remain to be understood.

**Fig. 5.**
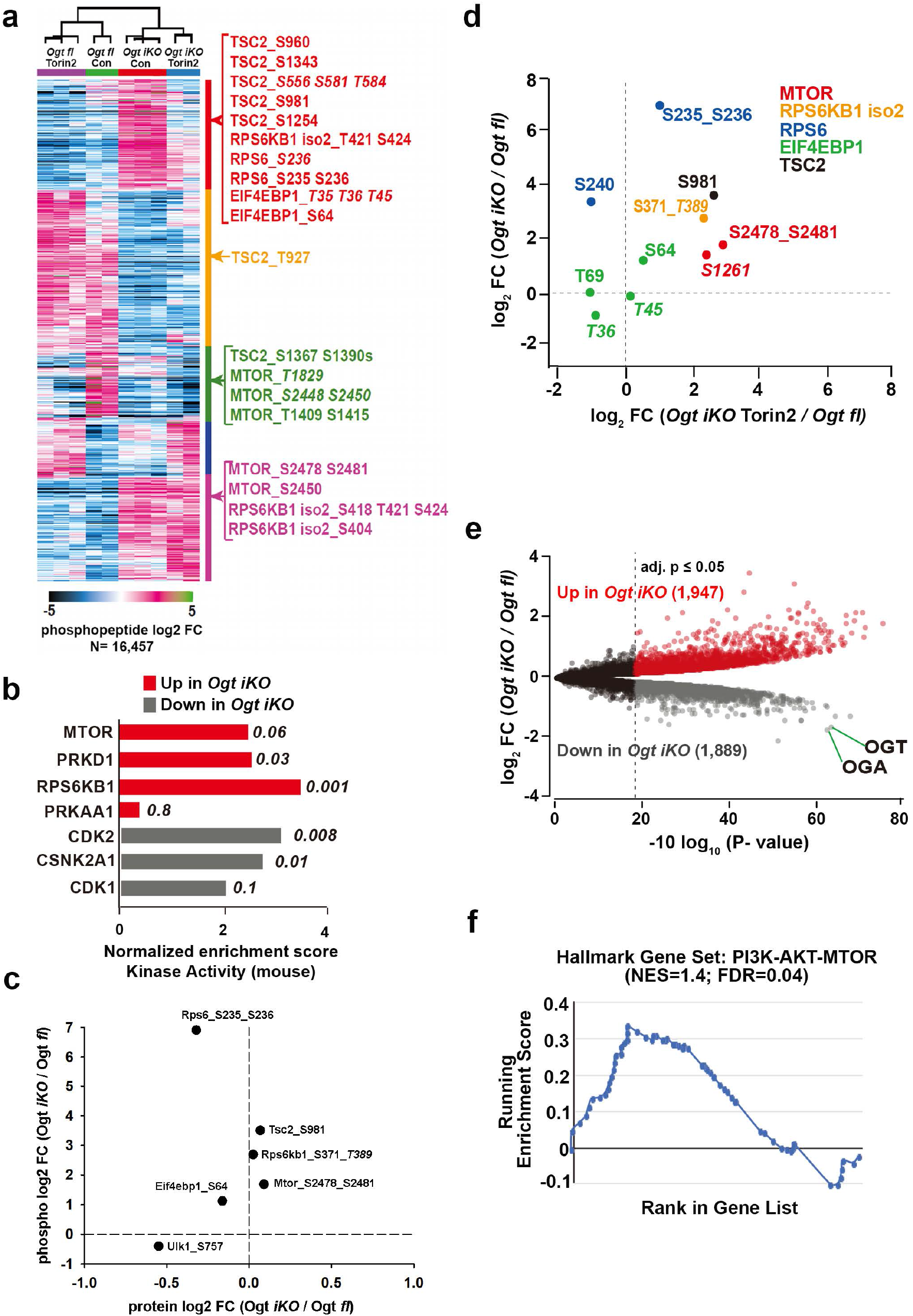
Extensive changes in global signaling and protein abundances in *Ogt iKO* mESCs. **a**, K-means clustering analysis of all regulated (B-H corrected adjusted p-value ≥ 0.05) phosphopeptides as determined by a moderated F-test. Example phosphopeptides for canonical mTOR pathway components are highlighted. Italicized phosphosite numbers indicate phosphosite assignment ambiguity. **b**, Normalized enrichment scores for kinase activity as determined by Post-translational modification-set enrichment analysis (PTM-SEA) of phospho-proteomics data performed in *Ogt fl* and *Ogt iKO* mESCs. Top three pathways in both directions, as well as their respective p-values are shown. PRKAA1 (AMPKA1) is included as a comparison. Only mouse-derived PTM sets were used for the analysis. **c**, Changes in phosphopeptide and protein levels of mTOR signaling components in *Ogt fl* and *Ogt iKO* mESCs as determined by quantitative proteomics. Italicized phosphosite numbers indicate phosphosite assignment ambiguity. **d**, Changes in phosphorylation levels on mTOR signaling components upon *Ogt* deletion alone (y-axis) and *Ogt* deletion in the presence of Torin2 (x-axis). Both are relative to *Ogt fl*. Italicized phosphosite numbers indicate phosphosite assignment ambiguity. “Iso 2” refers to isoform 2 of RPS6KB1. **e**, Volcano plot of differentially abundant proteins between *Ogt floxed* and *Ogt iKO* mESCs. B-H corrected p-values of less than or equal to 0.05 are highlighted. **f**, Leading edge analysis for proteome level-GSEA of proteins involved in the PI3K-AKT-MTOR pathway. Positive enrichment indicates up in *Ogt iKO*; negative, down in *Ogt iKO*. Normalized enrichment score and false discovery rate (FDR, q-value) are shown.

**Fig. 6.**
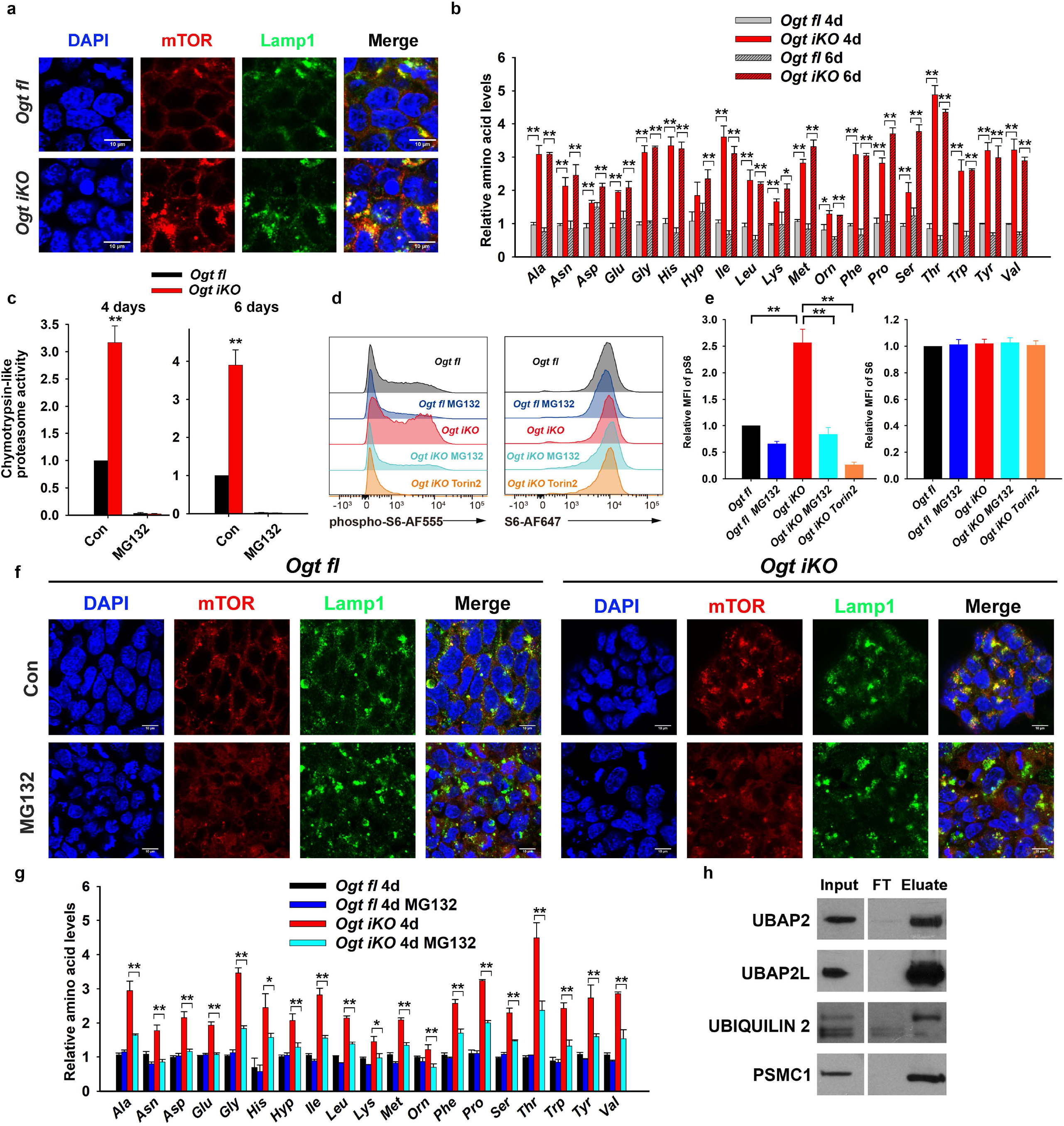
OGT deficiency promotes the translocation of mTOR by increasing proteasome activity. **a**, Immunohistochemistry of *Ogt fl* and *Ogt iKO* mESCs treated without or with 4-OHT respectively for 6 days. Cells were stained with antibodies against mTOR and Lamp1. Nucleus staining: DAPI (blue). Scale bar: 10 μm. **b**, Relative amino acid levels in *Ogt fl* and *Ogt iKO* mESCs treated without or with 4-OHT respectively for 4 or 6 days. Data are shown as mean ± SD (N=3). **c**, Chymotrypsin-like proteasome activity in *Ogt fl* and *Ogt iKO* mESCs treated without or with 4-OHT respectively for 4 or 6 days. Data are shown as mean ± SD (N=3). **d**, Flow cytometry analysis of phospho-S6 (*left panel*) and total S6 (*right panel*) ribosomal protein in *Ogt fl* and *Ogt iKO* mESCs treated without or with 4-OHT respectively for 6 days and then with or without 10 μM MG132 or 200 nM Torin2 for 2.5 hours. **e**, Relative mean fluorescence intensity (MFI) of phospho-S6 and S6 ribosomal protein in three independent experiments similar to that shown in (**d**). **f**, Immunohistochemistry of *Ogt fl* and *Ogt iKO* mESCs treated without or with 4-OHT respectively for 6 days and then with or without 10 μM MG132 for 2.5 hours. Cells were stained with antibodies against mTOR and Lamp1. Nucleus staining: DAPI (blue). Scale bar: 10 μm. **g**, Relative amino acid levels in *Ogt fl* and *Ogt iKO* mESCs treated without or with 4-OHT respectively for 4 days and then with or without 10 μM MG132 for 2.5 hours. **h**, Western blot of UBAP2, UBAP2L, Ubiquilin 2 and PSMC1 proteins after WGA pull down of cell lysates from *Ogt fl* mESCs. FT: Flow Through.

### mTOR is hyperactivated in OGT-deficient mESCs

Gene ontology analysis of the 115 candidate genes identified in our sgRNA screen highlighted six genes related to positive regulation of mTOR signaling (*Lamtor2, Lamtor3* and *Lamtor4, Rraga, Wdr24* and *Mios*) (**Fig. 4a**). Lamtor2, 3 and 4 are subunits of the Ragulator complex, which controls the activity of the Rag complex, a heterodimer of two guanine nucleotide-binding proteins, RagA (encoded by *Rraga*) and RagC (or alternately, RagB and RagD)^16^. Wdr24 and Mios are subunits of the GATOR2 complex, which indirectly senses amino acid levels and activate mTORC1 by inhibiting the GAP activity of GATOR1^13,14,16^. Since mTOR is an established regulator of mitochondrial function at both the transcriptional and translational level^18,19^, we examined mTOR activity in OGT-deficient cells. Analysis of mTOR activity at a single cell level by flow cytometry and immunostaining showed that the levels of phosphorylation of S6 ribosomal protein on serine 235/236, a reliable marker for mTOR activity, were strikingly increased in *Ogt iKO* mESCs, without a change in total S6 ribosomal protein levels; this increase was prevented by the ATP-competitive pan-mTOR inhibitor Torin2 (**Fig. 4b, c and Extended Data Fig. 7a, b**). Akt signaling, which is upstream of mTOR^27^, was also slightly increased in *Ogt iKO* mESCs (**Extended Data Fig. 7c, d)**. mTOR inhibitor Torin2 also restored the high oxygen consumption rate (**Fig. 4d** and **Extended Data Fig. 7e**), mitochondrial membrane potential and mitochondrial mass (**Extended Data Fig. 7f-i**) of *Ogt iKO* mESCs to nearly normal levels. At the transcriptional level, Torin2 also rescued the increased expression levels of genes encoded in mitochondrial DNA in *Ogt iKO* mESCs (**Extended Data Fig. 7j**). In contrast, the mitochondrial complex 1 inhibitor rotenone and an sgRNA targeting *Socs3*, which inhibited mitochondrial OXPHOS (**Fig. 3i**), did not affect mTOR hyperactivation in *OGT iKO* mESCs (**Extended Data Fig. 8a-d**). These data indicate that increased mitochondrial OXPHOS activity is largely downstream of mTOR hyperactivation. Consistent with our observation in *Ogt iKO* mESCs, inhibition of OGT catalytic activity by the OGT inhibitor OSMI-1^28^ significantly increased mTOR activity in control *Ogt fl* mESCs, as assessed by flow cytometry for phosphorylation of S6 ribosomal protein (**Extended Data Fig. 8e, f**).

Since mTOR signaling was strongly up-regulated upon *Ogt* deletion, we tested whether mTOR inhibition could rescue the viability of OGT-deficient mESCs. Treatment with the mTOR inhibitor Torin2, the mTORC1 inhibitor rapamycin or the AKT inhibitor AKTVIII partially rescued colony morphology and the decrease in mESC cell numbers observed in *Ogt iKO* cells (**Fig. 4e, f, Extended Data Fig. 9a**), despite the fact that *Ogt fl* cells showed a slight decrease in numbers upon treatment with these inhibitors. Treatment with Torin2 also rescued the expression of mRNAs encoding the pluripotency markers *Oct4* and *Nanog* in *Ogt iKO* cells (**Extended Data Fig. 9b**). Similarly, CRISPR-Cas9 mediated disruption of the genes encoding mTOR pathway components *Lamtor2, Lamtor4, Rraga* and *Wdr59* partially rescued cell numbers in *Ogt*-deleted cells, despite reducing cell numbers somewhat in *Ogt fl* mESCs (**Fig. 4g; Extended Data Fig. 9c, d**). Consistent with this observation, expression of sgRNA targeting *Lamtor2* inhibited hyperactivated mTOR signaling in *Ogt iKO* mESCs as judged by immunostaining for phosphorylation of S6 ribosomal protein (**Extended Data Fig. 9e**). mTOR pathway plays key roles in protein synthesis^29^ and has been reported to regulate mitochondrial function by regulation of protein synthesis^19^. Notably, cycloheximide, an inhibitor of protein synthesis that is toxic for control mESCs, rescued the decreased proliferation of *Ogt iKO* mESCs (**Extended Data Fig. 10a, b**). These results point to a functional link between OGT, mTOR activity and mitochondrial function that is essential for the proliferation and survival of OGT-deficient mESCs.

### Extensive changes in global signaling and protein abundance in *Ogt iKO* mESCs

To evaluate the global effects of mTOR activation in OGT-deficient cells, we performed quantitative proteomic and phosphoproteomic analysis of *Ogt fl* and *Ogt iKO* mESCs in the presence and absence of the mTOR inhibitor Torin2. Phosphoproteomics revealed extensive changes in signaling in mESCs lacking OGT (**Fig. 5a**). Assessment of kinase activity by post-translational modification-set enrichment analysis (PTM-SEA)^30^ showed significant upregulation of mTOR, RPS6KB1 (S6 kinase) and PKRD1 signaling pathways in OGT-deficient cells compared to *Ogt fl* cells (**Fig. 5b**). As a comparison, AMPKA1 (PRKAA1) kinase, which senses ATP levels in the cell, showed only minor activation upon *Ogt* deletion. Kinases associated with cell cycle progression, CDK1/2 and CSNK2A1 had diminished activity in OGT-deficient mESCs (**Fig. 5b**). Comparing changes in phosphorylation levels with changes in protein levels to assess phospho-site stoichiometry of canonical mTOR signaling components, we found that activating phosphorylations on mTOR and Rps6kb1 (an established target for phosphorylation by mTOR), were significantly upregulated upon *Ogt* deletion without changes in mTOR or Rps6kb1 protein levels (**Fig. 5c**). Ribosomal protein S6 (RPS6), a well-established target of S6K, showed a small though significant decrease in protein levels with a concomitant increase in phosphorylation levels at S235 and S236, indicating an increase in stoichiometry (**Fig. 5c; Extended Data Fig.11a)**. TSC2, a negative regulator of mTORC1 signaling, showed significant upregulation of its inhibitory phosphorylation site S981, also without changes in protein levels (**Fig. 5c**). The stoichiometry of phosphorylation (normalized to protein levels) of two downstream targets of mTOR – EIF4EBP1 at S64 and ULK1 at S757 – was also increased in *Ogt iKO* mESCs (**Fig. 5c; Extended Data Fig.11a)**.

Treatment of *Ogt iKO* cells with Torin2 decreased the abundance of ∼20% of the phosphopeptides dysregulated by *Ogt* deletion and subsequent mTOR activation (**Fig. 5a**). Amongst these were several phosphorylation sites known to be downstream of mTOR, including RPS6KB1, RPS6 and EIF4EBP1 (**Fig. 5d**). Torin2 did not alter a phosphorylation site, S1261, on mTOR that is responsive to amino acids^31^, suggesting that the ability of mTOR to sense changes in amino acid levels may remain intact in OGT-deficient cells. The proteome of *Ogt iKO* mESCs also showed major changes in protein levels (3,836 of the 8,470 measured) compared to *Ogt fl* cells, where OGT and OGA were among the most downregulated proteins upon *Ogt* deletion (**Fig. 5e**). Gene set enrichment analysis of the proteome-level data also showed increased expression of proteins in the PI3K-AKT-mTOR signaling pathway in *Ogt*-deficient mESCs (**Fig. 5f**). Consistent with the increased frequency of apoptotic cells in OGT-deficient cells (**Fig. 1g and Extended Data Fig.1c**), protein levels of the mitochondrial apoptosis markers CDKN1A (p21), CDKN2A (p16), CASP8, APAF1 and BAX were increased in *Ogt iKO* cells, and this increase could be rescued by the mTOR inhibitor Torin2 (**Extended Data Fig.11b**). Together these data show that *Ogt* deletion leads to the activation of mTOR/S6K signaling and extensive remodeling of the proteome and phosphoproteome.

### OGT deficiency promotes mTOR translocation by increasing proteasome activity

We investigated the mechanisms underlying aberrant mTOR activation in the absence of OGT. Notably, all six mTOR pathway hits from our CRISPR/Cas9 screen were related to mTOR translocation to the lysosomal membrane in response to amino acid sensing – mediated on the one hand by the Rag and Ragulator complexes (sgRNAs against *Rraga, Lamtor* 2-4), and on the other hand to amino acid sensing mediated by the GATOR2 complex (sgRNAs against *Wdr24* and *Mios*)^13,14^. Moreover, an amino acid-responsive phosphorylation site on mTOR was induced upon *Ogt* deletion and was not altered by Torin2 (**Fig. 5d**). We therefore assessed mTOR subcellular localization in *Ogt fl* and *Ogt iKO* mESCs by immunocytochemistry, and also monitored steady-state amino acid levels in these cells (**Fig. 6a, b**). In *Ogt fl* mESCs, mTOR staining was diffusely localized to the cytoplasm, with little overlap with the lysosomal marker LAMP1, whereas in *Ogt iKO* mESCs, mTOR showed perinuclear staining that overlapped considerably with LAMP1 (**Fig. 6a; Extended Data Fig. 12a**). sgRNA-mediated disruption of *Lamtor2* inhibited mTOR translocation to the lysosome in *Ogt iKO* mESCs (**Extended Data Fig. 12b**), confirming that mTOR was activated by translocation to the lysosome in the absence of OGT.

The lysosomal translocation and activation of mTOR correlated with a global increase in amino acid levels in *Ogt iKO* mESCs, apparent as early as 4 days or 6 days after 4-OHT treatment (**Fig. 6b**). Steady-state amino acid levels are controlled by the balance between protein synthesis, which depends on mRNA translation by ribosomes, and protein degradation, effected by proteasomes, lysosomes and autophagosomes. We focused on 26S proteasome activity, which was previously reported to be reversibly inhibited by OGT through direct *O*-GlcNAc modification of proteasome subunits including the ATPase Rpt2 (also known as Psmc1)^32^. Indeed, three major proteasome activities (chymotrypsin-like, trypsin-like, caspase-like) were all significantly increased in *Ogt iKO* compared to *Ogt fl* mESCs (**Fig. 6c; Extended Data Fig. 13a, b**), and this increase in activity was completely inhibited by treatment for short times (1.5 h) with 10 μM of the proteasome inhibitor MG132 (**Fig. 6c; Extended Data Fig. 13a, b**). Furthermore, an in-gel proteasome assay showed that 26S and 30S proteasome activities were significantly increased, while the activity of the 20S proteasome (which is present at low levels in ES cells) was slightly increased in *Ogt iKO* mESCs (**Extended Data Fig. 13c, d**). These data indicate that OGT inhibits the ATPase activity of the capped 26S and 30S proteasomes. However, the mitochondrial complex 1 inhibitor rotenone and the pan-mTOR inhibitor Torin2 could not block the increase of proteasome activity in *Ogt* iKO cells (**Extended Data Fig.13e**). Given that Torin2 significantly blocked the increase of mTOR activity (**Extended Data Fig. 13f, g**) and rotenone significantly blocked the increase of mitochondrial membrane potential in *Ogt iKO* cells (**Extended Data Fig. 13h, i**), our data suggest that mTOR hyperactivation occurs downstream of proteasome activity. To confirm the hypothesis that mTOR is hyperactivated at least partly because of increased proteasome activity, we measured mTOR hyperactivation in the presence or absence of proteasome inhibitors in control *Ogt fl* mESCs and *Ogt iKO* mESCs. Treatment of the cells for 2.5 h with MG132 or the structurally unrelated proteasome inhibitor Bortezomib significantly reduced phospho-S6 staining, assessed by flow cytometry and immunocytochemistry for phospho-S6, both in control cells and after *Ogt* deletion (**Fig. 6d, e; Extended Data Fig.14a-c**). The levels of total S6 ribosomal protein were unchanged under these conditions (**Fig. 6d, e; Extended Data Fig. 14a-c**). MG132 treatment also significantly reduced the lysosomal localization of mTOR in *Ogt iKO* mESCs, as indicated by the poor overlap of mTOR staining with the lysosome marker LAMP1 (**Fig. 6f; Extended Data Fig. 14d**), and partially rescued the aberrant increase of amino acids in *Ogt iKO* mESCs (**Fig. 6g**).

In addition to the direct O-GlcNAc modification of proteasome subunit Psmc1^32^, Ubiquilin-2, ubiquitin associated proteins Ubap2 (Ubiquitin-associated protein 2) and Ubap2l (Ubiquitin-associated protein 2-like) have been reported to be directly O-GlcNAcylated (**Supplementary Table 3**)^33,34^. Ubiquilin-2 links the proteasome and the polyubiquitin chains on targeted proteins through its N-terminal ubiquitin-like domain (UBL) and C-terminal ubiquitin-associated domain (UBA)^35^. Both Ubap2 and Ubap2l contain UBA domains and play important roles in the ubiquitin-proteasome pathway^36,37^. To corroborate these findings in mESCs, we precipitated whole cell lysates of *Ogt fl* mESCs with wheat germ agglutinin (WGA), a lectin that recognizes the *O*-GlcNAc modification, and immunoblotted for Pmsc1, Ubiquilin-2, Ubap2 and Ubap2l. These experiments confirmed previous findings (**Supplementary Table 3**)^34^ showing that Pmsc1/Rpt2, Ubiquilin-2, Ubap2 and Ubap2l were *O*-GlcNAc-modified in *Ogt fl* mESCs (**Fig. 6h**). In the case of Ubiquilin-2, we observed clear enrichment for a slowly-migrating, presumably *O*-GlcNAc-modified protein species. Together these data suggest that proteasomal subunit and ubiquitin associated proteins are *O*-GlcNAc modified by OGT; in the absence of OGT, high proteasome activity and consequent high intracellular amino acid levels result in aberrant mTOR activation and increased mitochondrial OXPHOS, thus causing decreased proliferation and viability of mESCs.

### Increased amino acid levels and increased mTOR activation are also observed in OGT-deficient T cells

Next, we asked if what we observed in mESCs is a general mechanism across mammalian cell types. To answer this question, we used CD8^+^ T cells isolated from male *Ogt*^*floxed*^*Cre-ERT2*^*+/KI*^ *Rosa26-LSL-YFP*^*+/KI*^ mice. At day 6 after 4-OHT treatment, the *O*-GlcNAc modification was completely eliminated in *Ogt iKO* CD8^+^ T cells as assessed by flow cytometry (**Extended Data Fig. 15a**). As expected, *Ogt iKO* CD8^+^ T cells showed a significant impairment of growth rate (**Extended Data Fig. 15b**; note logarithmic scale of Y-axis). Consistent with our results in mESCs, OGT deficiency in CD8^+^ T cells was associated with a significant increase in the levels of certain amino acids (**Extended Data Fig. 15c**), as well as with mTOR activation assessed by flow cytometry for phospho-S6 (**Extended Data Fig. 15d, e**). The levels of phosphorylation of S6 ribosomal protein on serine 235/236 were significantly increased in *Ogt iKO* CD8^+^ T cells while the levels of total S6 ribosomal protein were slightly decreased as analyzed by flow cytometry (**Extended Data Fig. 15d, e**). Treatment with the mTOR inhibitor Torin2 and the mitochondrial inhibitor Rotenone partially rescued the decrease of cell numbers in *Ogt iKO* CD8^+^ T cells, despite a decrease in cell numbers of control *Ogt fl* CD8^+^ T cells upon treatment with these inhibitors (**Extended Data Fig. 15f, g**). Together, these data suggest that OGT controls cell viability through the proteasome/ mTOR/mitochondrial axis in both embryonic stem cells and somatic cells.

## Discussion

The O-GlcNAc transferase OGT is highly conserved across species^1,38^ and is absolutely required for the proliferation and survival of mammalian cells^3,39,40^, but the reason for this requirement has been unclear^1^. To address this point, we performed an unbiased, genome-wide CRISPR-Cas9 “suppressor” screen in *Ogt fl* mESCs, in which the *Ogt* gene could be inducibly deleted by treatment with 4-OHT. The screen provided important insights, revealing that mitochondrial dysfunction is a major cause for the block in cell proliferation and eventual loss of cell viability induced by OGT deficiency. We traced this phenotype to mTOR hyperactivity, resulting from increased steady-state amino acid levels that in turn stemmed from increased proteasomal activity (**Extended Data Fig. 16**). An independent phosphoproteomic screen also emphasized the pronounced activation of the mTOR pathway in OGT-deficient mESCs. The data are generalizable to other cell types, since increased amino acid levels and increased mTOR activation are also observed in OGT-deficient CD8^+^ T cells. A useful outcome is that we have been able for the first time to obtain viable, proliferating, OGT-deficient mESCs by inhibiting mTOR hyperactivation with Torin2, or by depleting candidate gene products from the screen. The availability of cultured OGT-deficient cells will permit a comprehensive analysis of how OGT regulates diverse aspects of cellular function, in the absence of complications posed by cell cycle arrest and ensuing apoptosis.

Our study connects OGT deficiency directly to a signalling pathway that is initiated by proteasome hyperactivity and ends with arrested cell proliferation and eventual apoptotic cell death. The pathway is as follows: Loss of OGT activity → increased proteasome activity → increased intracellular amino acid levels → hyperactivation of mTOR → excessive increase of mitochondrial OXPHOS → cell proliferation arrest → loss of cell viability (**Extended Data Fig. 16**). Taking these steps in order, we have shown a clear connection between *Ogt* deletion, increased proteasome activity and increased amino acid levels in *Ogt iKO* mESC. Briefly, proteasome chymotrypsin-like, trypsin-like and caspase-like activities were all increased upon *Ogt* deletion, and in-gel proteasome activity assays showed that 20S, 26S and 30S proteasome activities were high in *Ogt iKO* mESC. Besides the proteasome subunit Psmc1 (formerly Rpt2)^32^, several ubiquitin-associated proteins, including UBAP2, UBA2L and UBIQUILIN2, were targets for *O*-GlcNAcylation by OGT. Intracellular amino acid levels were high in *Ogt iKO* compared to control *Ogt fl* mESC, and the increase in amino acid levels was blocked by the proteasome inhibitor MG132. Similarly, our studies in *Ogt iKO* mESC confirmed the connection between increased amino acid levels and increased mTOR activation, which occurs in multiple cell types^13^. Amino acids are essential activators for mTOR translocation to the lysosome, via a complex containing RAG GTPases, RAGULATOR and the vacuolar ATPase^13,41^. sgRNAs against RAGULATOR components partially rescued the arrest in cell proliferation observed in *Ogt iKO* mESC, and mTOR activation in *Ogt iKO* cells was blocked by MG132. We also observed a striking increase in mTOR translocation to the lysosomal membrane in *Ogt iKO* compared to control *Ogt fl* cells. Finally, mTOR regulates mitochondrial biogenesis in many cell types^18 13,19^, including ES cells as we show here. Briefly, the majority of genes encoded in mitochondrial DNA displayed increased levels of expression in *Ogt iKO* mESCs, and this increase was suppressed by the mTOR inhibitor Torin2. Torin2 also inhibited the increase in mitochondrial membrane potential, the increase in mitochondrial mass, and the increase of mitochondrial OXPHOS observed in *OGT iKO* mESCs; furthermore in both ES cells and T cells, Torin2 rescues the block in cell proliferation occurring after *Ogt* deletion.

There are also clear precedents in the literature demonstrating the connection between increased OXPHOS and loss of cell viability. Both ER stress induced by tunicamycin, and DNA damage induced by UV irradiation, led to BAX activation, mitochondrial membrane permeabilization and cell death in Rat-1 fibroblasts; these could all be inhibited by OXPHOS inhibitors, including the complex I inhibitor rotenone and two unrelated inhibitors of the F_0_F_1_-ATPase, oligomycin and aurovertin B)^25^. Similar data were obtained in MCF-7 breast cancer cells and HepG2 hepatocellular carcinoma^25^. Consistent with these findings, we show that OXPHOS is markedly increased in *Ogt iKO* compared to control *Ogt fl* mESC; moreover mass spectrometry of whole cell lysates indicates that the protein levels of the mitochondrial apoptosis markers BAX, CDKN1A (p21), CDKN2A (p16), CASP8 and APAF1 also show a clear increase, which is blocked by the mTOR inhibitor Torin2. Furthermore, the OXPHOS inhibitor Rotenone partially rescued the proliferation arrest of *Ogt iKO* mESCs in both mESC and CD8 T cells.

Our whole-genome CRISPR/Cas9 screen confirmed the direct connection of OGT deficiency to aberrations in mitochondrial function. The largest category of sgRNAs that rescued proliferation in *Ogt iKO* mESC was related to mitochondrial function, leading to the discovery that mitochondrial OXPHOS was markedly increased in *Ogt*-deleted compared to control mESC. Notably, sgRNA-mediated disruption of genes encoding three of the most prominent hits in our screen – Socs3, Xpo5, and Gtpbp8 – returned mitochondrial OXPHOS to almost normal levels while also rescuing the proliferation defect of *Ogt*-deficient cells. These proteins were not previously known to have specific mitochondrial functions. Gtpbp8, a member of the Obg family of P-loop-containing small G proteins, is annotated as mitochondrial^42^; a homologue, Gtpbp10, resides in the mitochondrial matrix and is required for proper maturation of the large subunit of the mitochondrial ribosome^43^. Gtpbp8 may have a related function, since several Mrpl proteins – components of the mitochondrial large ribosomal subunit – were also hits in our screen. Socs3 is a key negative regulator of Stat3, which is at least partially localized to mitochondria^44^. Xpo5 is an exportin that exports pre-miRNAs and tRNAs from the nucleus to the cytoplasm^45^. Further analysis of the functional roles of these three proteins in mitochondria, and the mechanism by which they suppress mitochondrial dysfunction in OGT-deficient cells, would be of considerable interest. Notably, rescue by the top sgRNA candidates identified in our whole-genome screen was incomplete; given that OGT has thousands of substrates, other mechanisms (in addition to mTOR hyperactivation and the associated mitochondrial dysfunction) might contribute to the defect in cell proliferation in *Ogt*-deficient cells.

Our screen also provided the first indication that *Ogt* gene deletion in mESCs was strongly associated with mTOR hyperactivation. Six of the candidate sgRNAs targeted upstream regulators on mTOR, prompting us to show increased translocation of mTOR to the lysosome and increased phosphorylation of the established mTOR targets S6 kinase and ribosomal protein S6 in *Ogt iKO* compared to *Ogt fl* mESC. All six mTOR-related candidates function in the pathway of mTOR amino acid sensing and mTOR activation at the lysosomal membrane^13^. Wdr59 is part of the GATOR2 complex, which communicates information about intracellular amino acid levels to mTORC1, while Lamtor2 and Lamtor4 are part of the Ragulator complex, which activates mTORC1 when RagA and RagC in the RagA/RagC heterodimer are GTP- and GDP-bound respectively. To determine whether mTOR hyperactivation increased mitochondrial oxygen consumption, or conversely whether increased OCR resulted in mTOR hyperactivation, we performed epistasis experiments. Rotenone is a well-known mitochondrial complex I inhibitor; Torin2 is a pan-mTOR inhibitor which competes with ATP for binding the mTOR kinase; rapamycin is an inhibitor of the mTOR-containing mTORC1 (Raptor) complex; and AKT VIII is a pan-AKT inhibitor that inhibits mTOR activation by its upstream kinase AKT. Torin2 not only rescued the cell proliferation defect, it also suppressed the increase in mitochondrial membrane potential, mitochondrial mass, oxygen consumption rate (OCR), and the increased expression of genes encoded in mitochondrial DNA. Despite its toxicity to normal cells, the mitochondrial complex 1 inhibitor rotenone partially rescued the proliferation defect of *Ogt*-deficient cells, as did rapamycin and the AKT inhibitor; notably, however, rotenone did not block mTOR hyperactivation in *Ogt iKO* cells. These data establish the direction of causality, showing that the mitochondrial dysfunction observed in OGT-deficient cells occurs largely downstream of mTOR hyperactivation. In other words, mTOR hyperactivation in OGT-deficient cells led to increased mitochondrial OXPHOS, which in turn led to arrested cell proliferation, not vice versa. A corollary is that the mTOR and mitochondrial OXPHOS pathways are causally connected, and are not independently responsible for cell viability downstream of OGT.

Many previous studies of the crosstalk between OGT and mTOR have focused on how mTOR signaling affects OGT function. In breast cancer cells, mTOR activation was associated with increased levels of OGT and *O*-GlcNAc modification^46^. In hepatic cancer cells, pharmacological inhibition of mTOR led to decreased OGT expression concomitant with a global reduction of *O*-GlcNAc modification^47^. Thus, in these settings, mTOR appears to activate OGT. Our focus here is in the opposite direction: i.e. how OGT affects mTOR activity. Because mTOR activity promotes cell proliferation^48^, and because of the dramatic decrease in cell proliferation in *Ogt iKO* cells, it was generally assumed that OGT deficiency would lead to decreased mTOR^49^. Consistent with this assumption, treatment of neurons with OGT inhibitors significantly downregulated mTOR activity^49^, suggesting that OGT might activate mTOR. Notably, however, nondividing cells such as neurons – unlike proliferating cells – can survive in the absence of OGT^50^, and given the differences in cellular context, the mechanism of crosstalk between OGT and mTOR has not been easy to decipher. However, a previous publication showed that mTOR activity (assessed by immunoblotting for phospho-S6K) was significantly increased in OGT-deficient CD8^+^ T cells, although the authors pursued a different finding, the downregulation of Myc protein in these cells^40^. Thus, our study is the first to show that complete *Ogt* gene deletion is (perhaps paradoxically) associated with mTOR hyperactivation, leading to increased mitochondrial OXPHOS, proliferation arrest and eventual loss of viability of Ogt *iKO* cells (**Extended Data Fig. 16**).

As mentioned above, all six mTOR-related candidates from our genome-wide screen functioned in the pathway of amino acid sensing by mTOR and its recruitment to and activation at the lysosomal membrane^13^. Amino acids are known to be essential activators of mTOR translocation to the lysosome^13^, and *Ogt iKO* mESCs displayed high amino acid levels and increased lysosomal localisation of mTOR compared to control *Ogt fl* mESCs. Steady-state amino acid levels are determined by the balance between protein synthesis and degradation^51-53^, and previous reports that the proteasome has an important role in controlling intracellular amino acid levels by degrading proteins to provide amino acids^51^, and that the *O*-GlcNAc modification suppresses proteasome activity *in vitro*^32^, led us to test the effects of OGT deficiency on proteasome function. We found that three different activities of the proteasome – chymotrypsin-like, trypsin-like and caspase-like activities – were all substantially increased upon *Ogt* deletion. The structurally unrelated proteasome inhibitors MG132 and Bortezomib, used for short times to reduce toxicity to normal cells, significantly reduced the increase in amino acid levels, as well as mTOR translocation and activity, in *Ogt iKO* cells. These data show unambiguously that OGT deficiency leads to increased proteasome activity, which in turn results in increased amino acids levels within the cell. Neither the mitochondrial complex 1 inhibitor rotenone nor the mTOR inhibitor Torin2 blocked the increase of proteasome activity and amino acid levels in *Ogt iKO* cells, indicating that proteasome activity is upstream of mTOR hyperactivation and increased mitochondrial OXPHOS (**Extended Data Fig. 16**).

## Acknowledgements

We thank Dr. Joyce Chen and Dr. Anand Balasubramani for generating *Ogt*^*floxed*^ *Cre-ERT2*^*KI/KI*^ mice and *Ogt*^*floxed*^ *Rosa26-LSL-YFP*^*KI/KI*^ mice; Dr. Mohit Jain (UCSD), Dr. Lucas Sullivan (Fred Hutchinson Cancer Research Center), and members of the Rao laboratory for suggestions and discussions. We thank Dr. Natasha Zachara from Johns Hopkins University School of Medicine for providing *Ogt floxed* mice; Dr. Ronald M. Evans (Salk Institute) for providing wildtype OGT plasmid; C. Kim, D. Hinz, C. Dillingham, M. Haynes and S. Ellis at the La Jolla Institute Flow Cytometry facility for help with cell sorting experiments; J. Day, S. Alarcon, H. Dose, K. Tanguay and A. Hernandez of the La Jolla Institute Sequencing facility for help with next-generation sequencing; and Olga Zagnitko at Cancer Metabolism Core at Sanford Burnham Prebys Medical Discovery Institute for assistance with GC/MS analyses. This work was supported by National Institutes of Health (NIH) R01 grant R35 CA210043 (to A.R.). FACSAria II Cell Sorter was acquired through the Shared Instrumentation Grant (SIG) Program S10 RR027366 and Hiseq 2500 was funded by S10OD016262. The Sanford Burnham Prebys Cancer Metabolism Core is supported by NCI Cancer Center Support Grant P30 CA030199. X.L. was supported by a postdoctoral Fellowship from CIRM UCSD Interdisciplinary Stem Cell Research & Training Grant II (TG2-01154). H.S. was supported by Pew Latin American Fellows Program from the Pew Charitable Trusts.

## Author contributions

A.R. and X.L. conceived the project. X.L. and X.Y. performed experiments, analyzed and interpreted the results. X.L. prepared samples for seahorse, GC/MS, proteomic and phosphoproteomic experiments. H.S. assisted with experiments. D.A.S. performed the seahorse experiments and GC/MS profiling of amino acids. R.A.B and S.A.M. performed the proteomic and phosphoproteomic experiments, analyzed and interpreted the data. S.A.C supervised the proteomic and phosphoproteomic experiments. A.R. supervised project planning and execution. X.L., X.Y., S.A.M. and A.R. wrote the manuscript, and all authors proofread the manuscript and provided editorial input.

## Materials and Methods

### Mice

*Ogt floxed* mice^3,39^ backcrossed to C57BL/6 mice for 12 generations were obtained from Dr. Natasha Zachara (Johns Hopkins). B6.129-*Gt(ROSA)26Sor*^*tm1(cre/ERT2)Tyj*^/J strain (stock number: 008463; *Cre/ERT2* is a fusion protein of Cre with a modified estrogen receptor, which enters the nucleus only after it binds 4-hydroxytamoxifen) and B6.129×1-*Gt(ROSA)26Sor*^*tm1(EYFP)Cos*^/J strain (stock number: 006148; LSL: loxP-STOP-loxP cassette contains a strong transcriptional STOP site flanked by loxP sites, which turns on EYFP expression upon Cre-mediated deletion of *loxP* sites) were purchased from the Jackson Laboratory. All mice were on the B6 background and maintained in a specific pathogen-free animal facility in the La Jolla Institute for Immunology. All animal procedures were reviewed and approved by the Institutional Animal Care and Use Committee of the La Jolla Institute for Immunology and were conducted in accordance with institutional guidelines.

### Generation and culture of mouse embryonic stem cells

*Ogt*^*floxed*^ *Cre-ERT2*^*KI/KI*^ mice were crossed with *Ogt*^*floxed*^ *Rosa26-YFP*^*KI/KI*^ mice to obtain *Ogt*^*floxed*^*Cre-ERT2*^*+/KI*^ *Rosa26-LSL-YFP*^*+/KI*^ blastocysts (embryonic day 3.5) to generate male and female *Ogt floxed* mESC lines. Ogt deletion was then induced by the addition of 1 μM 4-hydroxytamoxifen (4-OHT) (TOCRIS, catalog number: 3412). Mouse ESCs were maintained on mitomycin C-treated mouse embryonic fibroblasts (MEFs; feeder cells) with LIF in Knockout DMEM medium (ThermoFisher Scientific, catalog number: 10829018) supplemented with 15% KOSR (KnockOut Serum Replacement, ThermoFisher Scientific, catalog number: 10828028), 2 mM L-Glutamine, 1 X MEM Non-Essential Amino Acids, 50 μM β-Mercaptoethanol. Cell numbers were counted by flow cytometry on a BD Accuri C6 (BD Biosciences).

### Plasmids

Wildtype *Ogt* plasmid was kindly provided by Dr. Ronald M. Evans (Salk Institute). Wildtype *Ogt* was subcloned into a lentiviral vector pLV-EF1a-IRES-Puro (Addgene #85132).

### Lentiviral packaging and transduction

Lentiviral packaging was performed in HEK293T cells using Lipofectamine 2000 (ThermoFisher Scientific) according to the manufacturer’s protocol. Briefly, pCMV-VSVG (Addgene, #8454), psPAX2 (Addgene, #12260) and lentiGuide plasmids were transfected at the ratio of 1:2:2. Lentiviral supernatant was harvested at 48 hr post-transfection and used for mESC infection.

### CRISPR/Cas9 Screen

An *Ogt*^*floxed*^*Cre-ERT2*^*+/KI*^ *Rosa26-LSL-YFP*^*+/KI*^ mESC cell line stably expressing Cas9 was generated by lentiviral transduction of the parental cells with lentiCas9-Blast (Addgene #52962). CRISPR sgRNA library lentiGuide-Puro-Brie (Addgene, #73633), which contains a pool of 78,637 sgRNAs targeting 19,647 genes was transduced into mESCs with MOI < 0.3 and an average of 400x coverage. After puromycin selection for 7 days to obtain cells stably expressing the sgRNAs, the cells were split into two groups, an untreated (*Ogt fl*) group that served as the control, and a second group treated with 4-OHT for 6 days to induce Ogt deletion (*Ogt iKO*). On day 13, YFP-negative cells from the control *Ogt fl* group and YFP^+^ cells from the *Ogt iKO* group were sorted and used for library preparation for next generation deep sequencing.

### Library Preparation, sequencing and data analysis

Genomic DNA was isolated using Quick-gDNA MidiPrep kit (Zymo Research, D3100) according to the manufacturer’s protocol. Library preparation was performed as described in (Feng Zhang, 2017, Nat Prot). sgRNA inserts were PCR-amplified using NEBNext High Fidelity PCR Master Mix (NEB, M0541). The resulting PCR amplicons were purified using Ampure XP Beads and sequenced using the HiSeq 2500 system (Illumina) to assess changes in abundance of sgRNAs between *Ogt fl* and *Ogt iKO* groups. The sequencing data were analyzed using the PinAPL-Py platform^22^. SigmaFC is a gene score by taking the sum of the log fold-changes of all sgRNAs targeting that gene, multiplying by the number of sgRNAs that reached statistically significant enrichment. SigmaFC is a gene score by taking the sum of the log fold-changes of all sgRNAs targeting that gene, multiplying by the number of sgRNAs that reached statistically significant enrichment.

### Mitochondria staining

mESCs were incubated with 100 nM MitoTracker red CMXRos (Thermofisher, M7512) or MitoTracker deep red (Thermofisher, M22426) or Mtphagy Dye (Dojindo, MT02-10) for 30 min at 37 °C. Cells were then subjected to flow cytometry using a FACS flow cytometer (BD Biosciences).

### Reactive oxygen species (ROS) measurement

ROS generation was determined by CellROX Deep Red (Thermofisher, C10422). The fluorescence intensity of CellROX Deep Red reflects the ROS levels. mESCs were incubated with 500 nM CellROX Deep Red for 30 min at 37 °C. Cells were then subjected to flow cytometry using a FACS flow cytometer (BD Biosciences).

### Cell viability measurement

Cells were stained with Fixable Viability Dye eFluor 780 (1:1000) (Thermofisher, 65-0865-18) for 30 min at 37 °C. Cells were then subjected to flow cytometry using a FACS flow cytometer (BD Biosciences).

### Seahorse Mito Stress assay and amino acid measurement

A Seahorse XFe24 Bioanalyzer (Agilent) was used to determine OCR (oxygen consumption rate). 100,000 mESCs per well were seeded in a Seahorse XFe24 cell culture plate precoated with Matrigel Matrix (Corning) one day prior to the assay. OCR was measured at baseline and after sequential addition of (i) oligomycin (2 μM; an ATP synthase inhibitor); (ii) carbonyl cyanide-4-(trifluoromethoxy) phenylhydrazone (FCCP) (1 μM; a mitochondrial uncoupler); (iii) rotenone (1 μM; a complex I inhibitor) and antimycin A (1 μM; a complex III inhibitor).

### Metabolite and amino acid measurement

1M mESC samples were collected in 50% methanol with 20 μM L-Norvaline. The levels of metabolites and amino acids were profiled by gas chromatography/mass spectrometry (GC/MS) at Cancer Metabolism Core, Sanford Burnham Prebys Medical Discovery Institute using methods described in reference^54^. Arginine is unstable in this analytical procedure and is converted to ornithine.

### Proteasome activity measurement

Three major proteasome activities (chymotrypsin-like trypsin-like, caspase-like) were measured using the Proteasome-Glo Cell-Based Assay Kit (G1180, Promega) as described by the manufacturer. Briefly, mESCs were trypsinized and washed with culture medium, 10,000 cells per well were seeded into a 96-well plate and incubated with or without 10 μM MG132 for 90 min at 37°C, 5% CO_2_. The plate was then allowed to equilibrate to room temperature for 15 min before the addition of 100 μL proteasome Glo reagent containing the bioluminescent substrate. The luminescent signals produced upon cleavage were measured using a SpectraMax Luminometer.

### In-gel proteasome activity assay

In-gel proteasome activity assay was performed as described^55^. Briefly, native protein extracts were prepared and resolved on native gel. The gel was incubate for 30 min with the reaction buffer at 37 °C in the dark box. After incubation, gel was transferred from the tray to Gel Doc (Bio-Rad) for analysis.

### Quantitative real-time PCR (qRT-PCR)

Quantitative real-time PCR was performed using Universal SYBR Green Master Mix (Roche) and analyzed using a Step One Plus real-time PCR system (Applied Biosystems) according to the manufacturer’s instructions and the data were normalized for *Gapdh* expression. The primers used for qRT-PCR are listed in **Supplementary Table 4**.

### Immunohistochemistry

Immunohistochemistry was performed as described previously^56^ with the primary antibodies (Clone name, conjugated fluorescence, dilution, manufacturer and catalog number shown in brackets): anti-*O*-Linked N-Acetylglucosamine antibody (RL2, Alexa Fluor 647, 1:200, Abcam, ab201994); Phospho-S6 Ribosomal Protein (Ser235/236) (D57.2.2E, Alexa Fluor 555, 1:200, Cell Signaling Technology, 39855); S6 Ribosomal Protein (54D2, Alexa Fluor 647; 1:200, Cell Signaling Technology, 55485); Lamp1 (1D4B, unconjugated, 1:50, Developmental Studies Hybridoma Bank, 1D4B); mTOR (7C10, unconjugated, 1:200, Cell Signaling Technology, 2983S); Phospho-Akt (Ser473) (D9E, Alexa Fluor 647, 1:100, Cell Signaling Technology, 4075). The colocalization correlation coefficients (Pearson’s correlation) between mTOR and Lamp1 were analyzed using software ImageJ2 with the plugin Coloc 2.

### Flow cytometry

mES cells were trypsinized into single cells and used for staining or cell sorting. The antibodies used for the staining are as following (Clone name, conjugated fluorescence, dilution, manufacturer and catalog number shown in brackets): Phospho-S6 Ribosomal Protein (Ser235/236) (D57.2.2E, Alexa Fluor 555, 1:200, Cell Signaling, 39855); S6 Ribosomal Protein (54D2, Alexa Fluor 647; 1:200, Cell Signaling, 55485); anti-*O*-Linked N-Acetylglucosamine antibody (RL2, Alexa Fluor 647, 1:200, Abcam, ab201994).

### Pulldown of O-GlcNAcylated proteins in cell lysates with wheatgerm agglutinin (WGA)

WGA pull down was performed using Pierce™ Glycoprotein Isolation Kit, WGA (Thermo Scientific) as described in the manufacturer’s protocol. Briefly, 800 μg of total cell lysates prepared with RIPA buffer were added to 200 μl of WGA lectin resin in a total volume of 500 μl bind buffer and incubated with mixing for 20 minutes at room temperature. The beads were washed four times in wash buffer; *O*-GlcNAcylated proteins were eluted from the resin at 95 °C, and the resulting eluent and the first flow-through from the column were analyzed by Western blotting with antibodies against PSMC1 (Proteintech, 11196-1-AP), Ubiquilin 2 (Novus Biologicals, H00029978-M03), UBAP2 (Bethyl Laboratories Inc, A304-627A), UBAP2L (Bethyl Laboratories Inc, A300-533A).

### Proteomics and Phosphoproteomics

mESC pellets were pelleted, snap-frozen with liquid nitrogen, and stored at −80°C until prepared for mass spectrometric analysis. Cell pellets were processed and analyzed as previously described with minor modifications^57,58^. Samples were analyzed on a Q-Exactive HF-X quadrupole Orbitrap mass spectrometer (Thermo Fisher Scientific).

Mass spectrometric data was analyzed using Spectrum Mill (Agilent) searching a Uniprot mouse database (12/28/2017) with 47,069 entries including common laboratory contaminants. All phosphosite assignments with less than 99% localization confidence are italicized and reported by Spectrum Mill without modification. TMT 11 ratios were generated using the median of all channels. Biological replicates with a Pearson’s r correlation of less than 0.2 were excluded. This was applied to one sample in the phosphoproteomics analysis, which is less stable than proteome changes^58^. The moderated T-test (pairwise comparisons) and moderated F-test (comparisons across all samples) were performed using Limma. GSEA was performed as previously described using only Hallmark gene sets^59^. PTM-SEA was performed as previously described, using only the mouse database^30^.

### CD8^+^ T cell isolation and activation

CD8^+^ T cells were isolated from spleen and lymph nodes of *Ogt*^*floxed*^*Cre-ERT2*^*+/KI*^ *Rosa26-LSL-YFP*^*+/KI*^ mice using Dynabeads Untouched Mouse CD8 Cells Kit (ThermoFischer, 11417D), and activated with plate-bound anti-CD3 (clone 2C11, BioXcell) and anti-CD28 (clone 37.51, BioXcell) antibodies at 1 μg/ml in the presence of 100 U/ml recombinant human IL-2 (rhIL-2). 4-OHT (1 μM) was added to induce *Ogt* deletion.

### Statistical Analysis

All values are shown as means ± SD. To determine the significance between groups, comparison was made using Student’s t test. For all statistical tests, the 0.05 confidence level was considered statistically significant. In all figures, * denotes p<0.05 and ** denotes p<0.01 in an unpaired Student’s t-test.

### Data Availability

The authors declare that all data supporting this study are available within the article, extended data figures and supplementary tables or from the corresponding authors upon reasonable request. A reporting summary for this article is available as a supplementary information file.

