## Supplementary Figures for "OGT controls mammalian cell viability by regulating the proteasome/mTOR/mitochondrial axis"

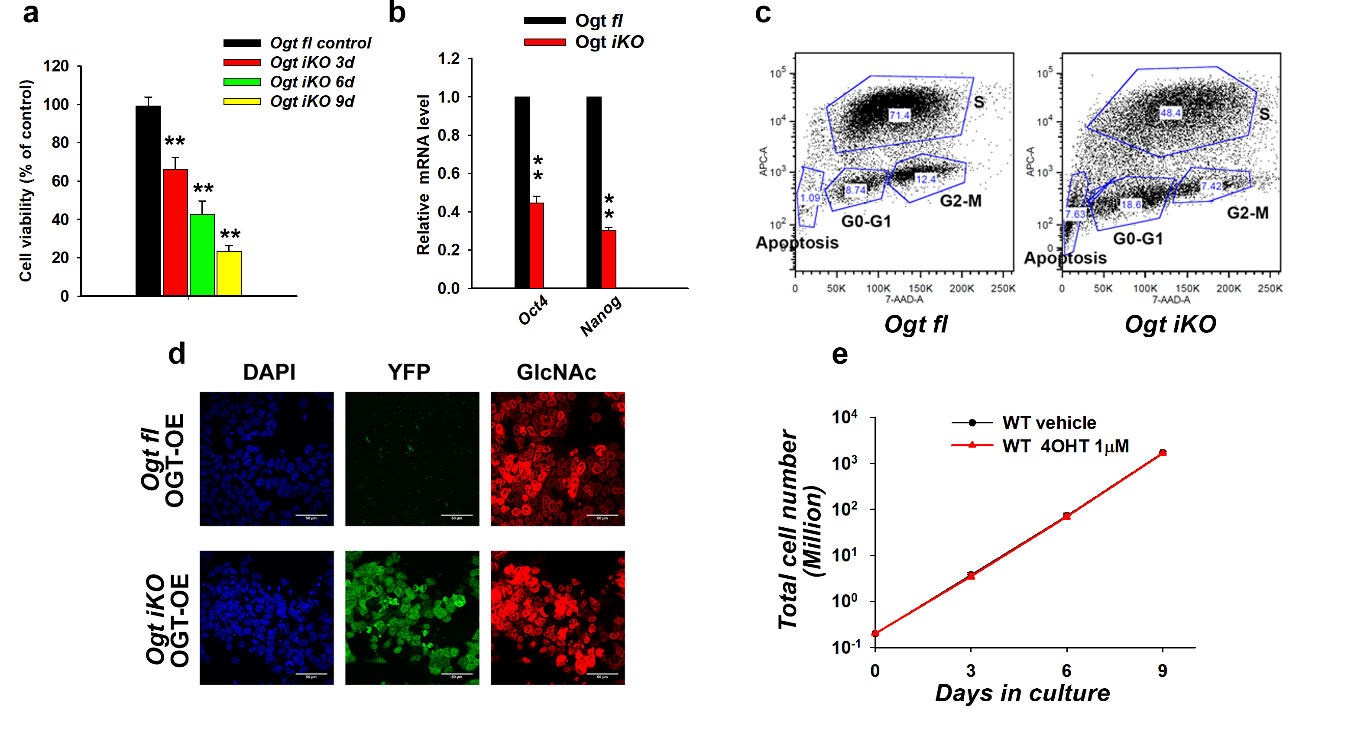

**Extended Data Fig. 1** | **Inducible deletion of *Ogt* impairs cell viability in mESCs.**

**a,** Cell viability of *Ogt fl* control and *Ogt iKO* mESCs 3, 6, and 9 days after 4-OHT treatment. Dead cells were labelled with fixable viability dye eFluor 780.

**b,** Quantitative real-time PCR (qRT-PCR) analysis of *Oct4 and Nanog* transcript levels in *Ogt fl* and *Ogt iKO* mESCs 6 days after 4-OHT treatment. The expression level of mRNA is shown relative to the level in control mESCs. Data are shown as mean ± SD (N=3).

**c,** Distribution of *Ogt fl* and *Ogt iKO* mESCs 6 days after 4-OHT treatment in different phases of the cell cycle. Representative dot blots showing the number of cells in S phase and DNA content by APC-BrdU and 7-AAD staining respectively.

**d,** Immunohistochemistry of *Ogt fl* and *Ogt iKO* mESC reconstituted with WT OGT treated with 4-OHT for 6 days, using antibodies against *O*-GlcNAc. Nucleus staining: DAPI (blue). Scale bar: 10 μm.

**e,** The cumulative growth curves of WT mESCs treated with or without 4-OHT are identical. Data are shown as mean ± SD (N=3).

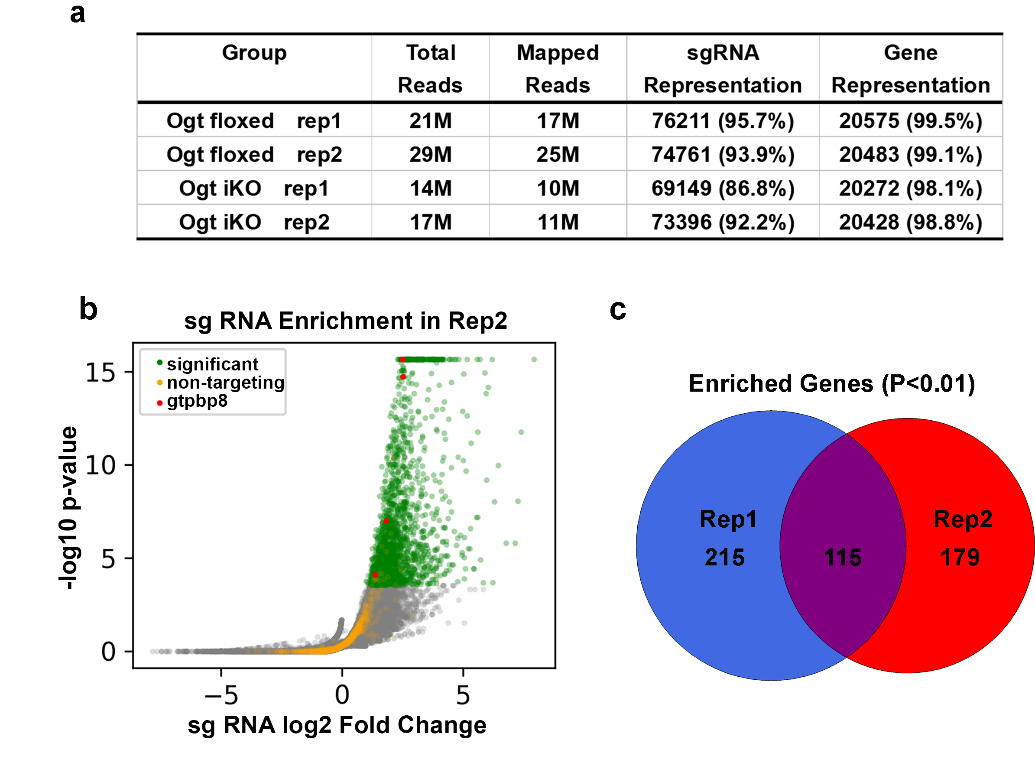
**Extended Data Fig. 2** | **Genome-wide CRISPR-Cas9 screen to identify key regulators of cell viability in OGT-deficient mESCs.**

**a.** Numbers of total and mapped reads for sgRNAs recovered from 2 replicates each of *Ogt fl* and *Ogt iKO* cells in the genome-wide CRISPR-Cas9 screen. The representation of sgRNAs and their corresponding genes is shown.

**b.** Volcano plot of p-value vs fold enrichment for each sgRNA in Replicate 2. Significantly enriched sgRNAs are shown in green and non-targeting control sgRNAs in orange. All four sgRNAs for *Gtpbp8* were significantly enriched in the screen (red dots).

**c.** Venn diagram showing the overlap of hits targeted by sgRNAs that were enriched in both independent CRISPR/Cas9 screens. A total of 115 hits scored highly in both replicates.

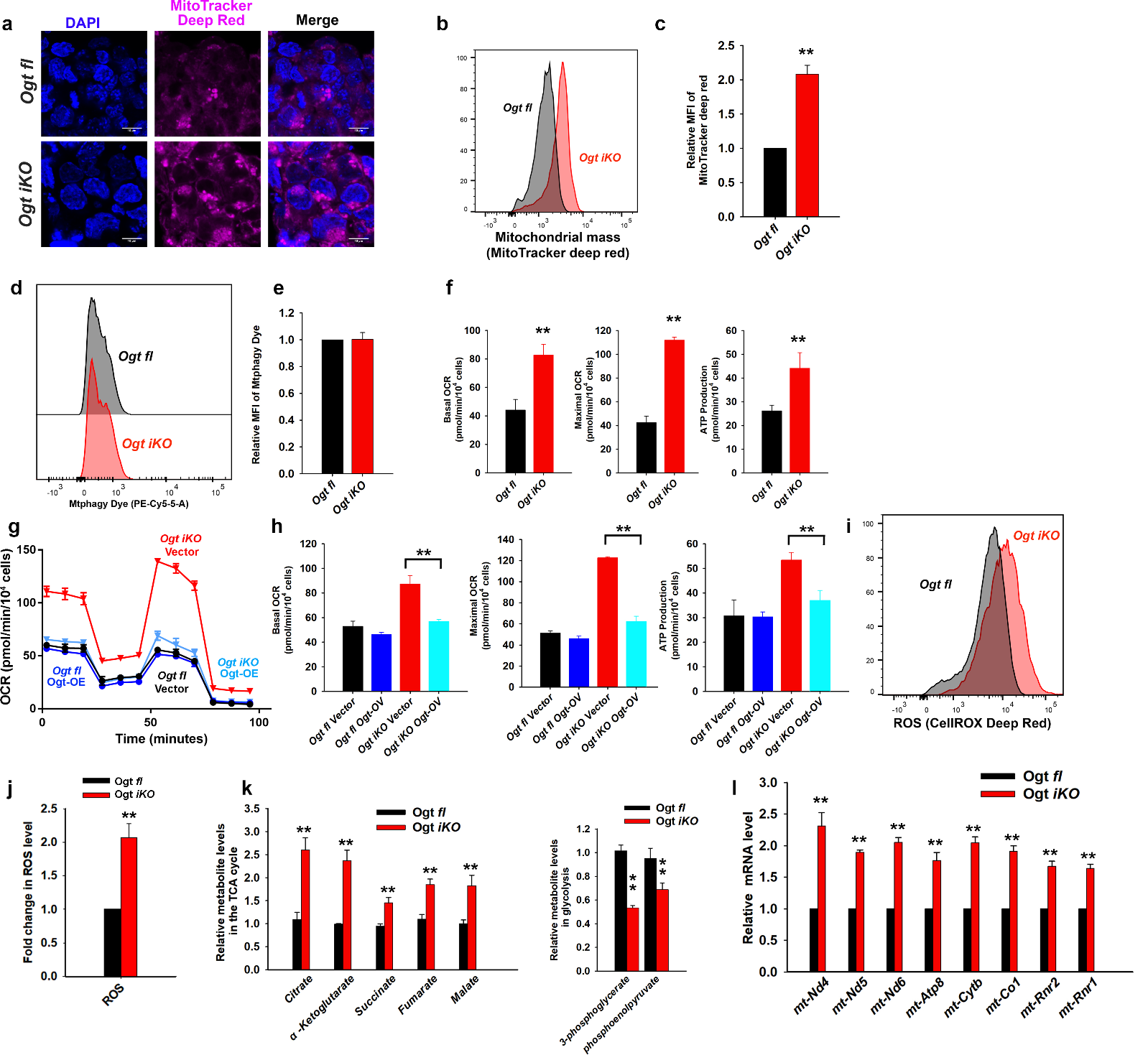

**Extended Data Fig. 3** | **OGT deficiency results in mitochondrial dysfunction.**

**a,** Fluorescence images of *Ogt fl* and *Ogt iKO* mESCs 6 days after 4-OHT treatment. Cells were stained with MitoTracker Deep red. Nucleus staining: DAPI (blue). Scale bar: 10 μm.

**b,** Representative histogram of MitoTracker deep red fluorescence in *Ogt fl* and *Ogt iKO* mESCs 6 days after 4-OHT treatment.

**c,** Relative MFI (mean fluorescent intensity) of MitoTracker deep red shown in **b**. Data are shown as mean ± SD (N=3).

**d,** Representative histogram of Mtphagy dye fluorescence in *Ogt fl* and *Ogt iKO* mESCs 6 days after 4-OHT treatment.

**e,** Relative MFI (mean fluorescent intensity) of Mtphagy dye shown in **d**. Data are shown as mean ± SD (N=3).

**f,** Basal OCR, maximal OCR, and ATP production in *Ogt fl* and *Ogt iKO* mESCs treated without or with 4-OHT respectively for 6 days. Data are shown as mean ± SD (N=3).

**g,** Analysis of oxygen consumption rate (OCR) using Seahorse XFe24 in *Ogt fl* mESCs stably expressing with empty vector or wildtype OGT treated with or without 4-OHT respectively for 6 days. Data are shown as mean ± SD (N=3).

**h,** Basal OCR, maximal OCR and ATP production in *Ogt fl* mESCs stably expressing empty vector or wildtype OGT and treated with or without 4-OHT respectively for 6 days. Data are shown as mean ± SD (N=3).

**i,** Representative histogram of ROS (cellROX deep red) in *Ogt fl* and *Ogt iKO* mESCs 6 days after 4-OHT treatment.

**j,** Relative MFI (mean fluorescent intensity) of ROS (cellROX deep red) shown in **i**. Data are shown as mean ± SD (N=3).

**k,** Relative metabolite levels of TCA cycle and glycolysis metabolites in *Ogt fl* and *Ogt iKO* mESCs 6 days after 4-OHT treatment.

**l,** qRT-PCR analysis of mitochondrial DNA-encoded gene transcript levels in control *Ogt fl* and *Ogt iKO* mESCs 6 days after 4-OHT treatment. The expression level of mRNA is shown relative to the level in control mESCs. Data are shown as mean ± SD (N=3).

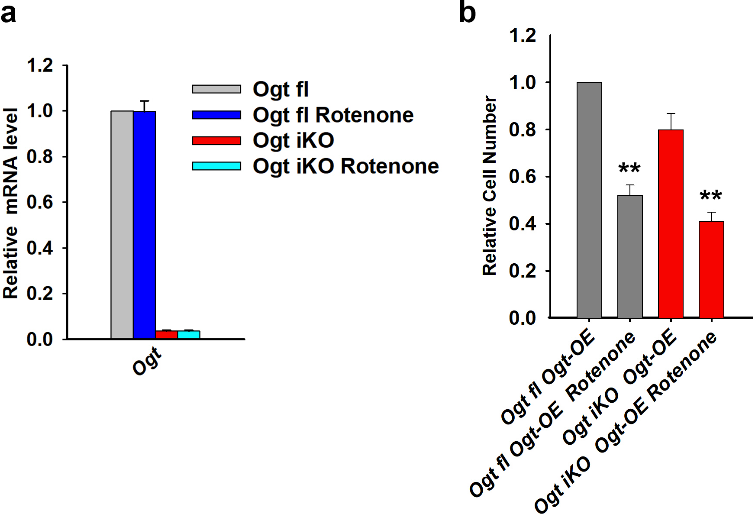

**Extended Data Fig. 4 | Rescue experiment by mitochondrial Complex I inhibitor rotenone.**

**a,** qRT-PCR analysis of *Ogt* transcript levels in *Ogt fl* and *Ogt iKO* mESCs treated with or without the mitochondrial Complex I inhibitor rotenone (75 nM). The expression level of *Ogt* mRNA is shown relative to the level in control mESCs. Data are shown as mean ± SD (N=3).

**b.** Relative cell numbers of *Ogt fl* mESCs stably expressing with wildtype OGT treated without or with 4-OHT respectively and with or without the mitochondrial Complex I inhibitor rotenone (75 nM) for 8 days. Data are shown as mean ± SD (N=3).

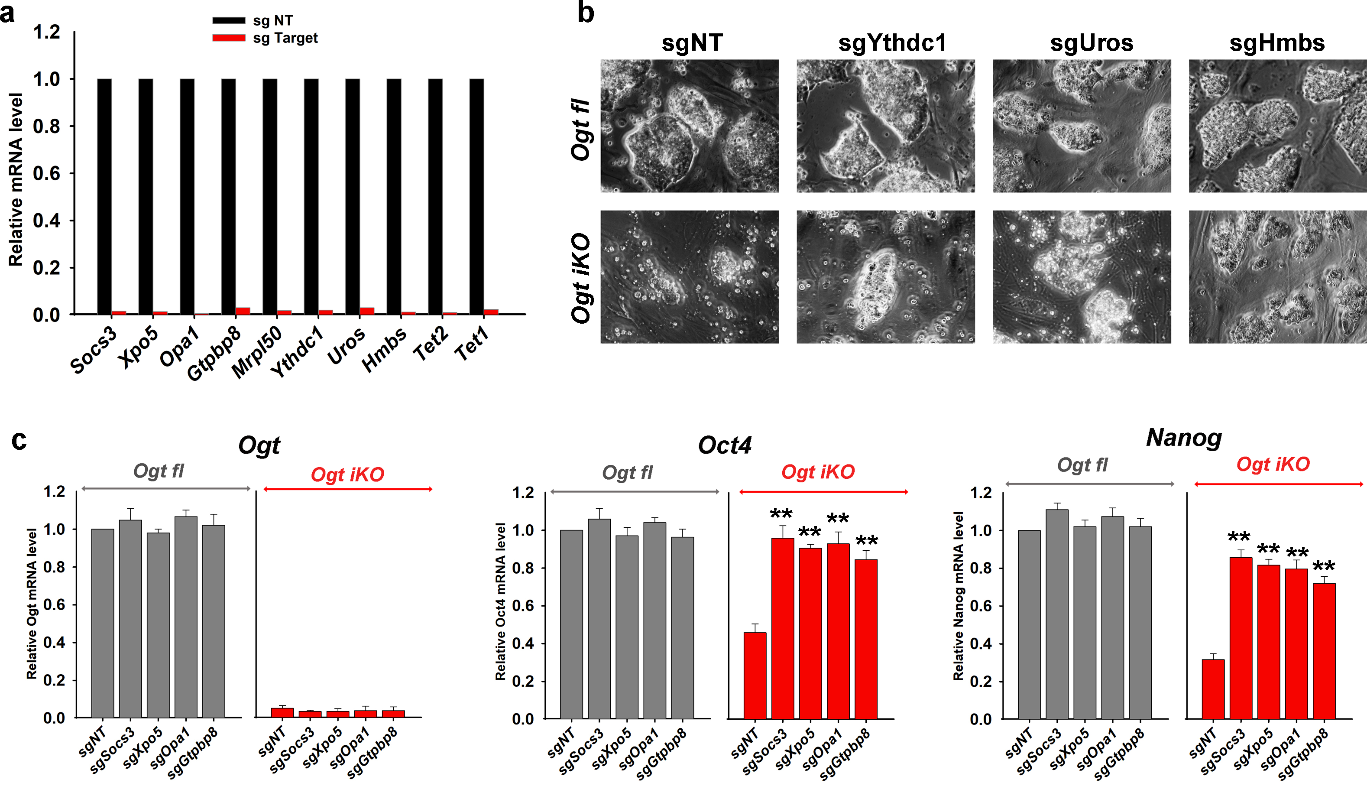

**Extended Data Fig. 5 | Rescue experiment by sgRNA identified in** **CRISPR-Cas9 screen.**

**a,** Efficient sgRNA-mediated depletion of mRNAs for the indicated genes in *Ogt fl* mESCs after lentiviral transduction of individual sgRNAs followed by puromycin selection for 7 days.

**b,** Phase contrast images of *Ogt fl* and *Ogt iKO* mESC colonies expressing non-targeting sgRNA (sgNT) or sgRNAs targeting the indicated genes (*Ythdc1*, *Uros*, *Hmbs*) after lentiviral sgRNA transduction followed by treatment with 4-OHT or vehicle for 8 days.

**c,** qRT-PCR analysis of *Ogt*, *Oct4 and Nanog* transcript levels in *Ogt fl* and *Ogt iKO* mESCs expressing non-targeting sgRNA (sgNT) or sgRNAs targeting the indicated genes (*Socs3*, *Xpo5*, *Opa1* and *Gtpbp8*) and treated without or with 4-OHT respectively for 6 days. The expression level of mRNA is shown relative to the level in control mESCs. Data are shown as mean ± SD (N=3).

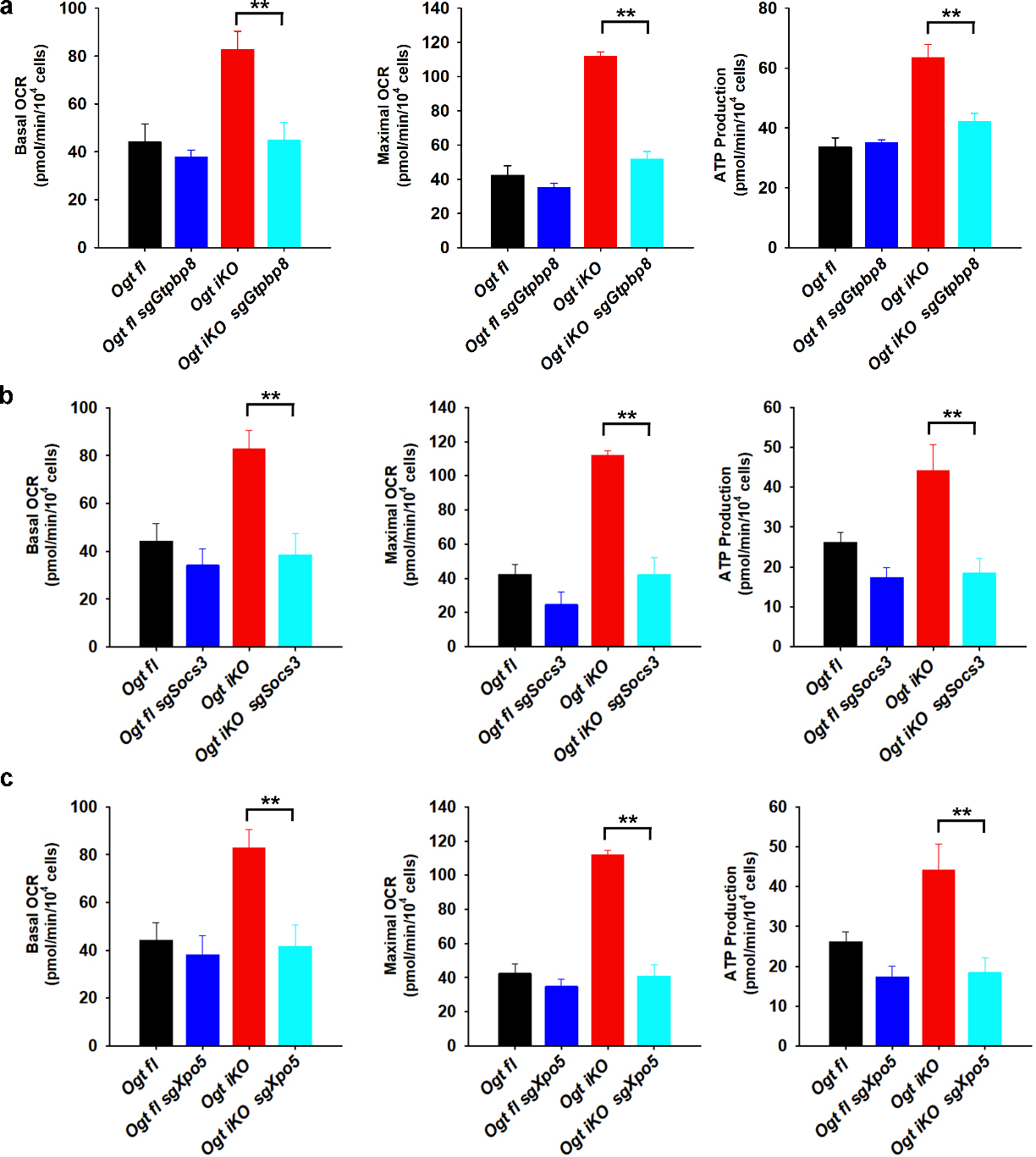

**Extended Data Fig. 6 | Mitochondrial Respiratory Profiles of sgRNA-expressing mESCs.**

**a-c,** The basal OCR, maximal OCR and ATP production in *Ogt fl* and *Ogt iKO* mESCs expressing sgRNAs targeting *Gtpbp8* (**a**), *Socs3* (**b**) and *Xpo5* (**c**) and treated without or with 4-OHT respectively for 6 days. Data are shown as mean ± SD (N=3).

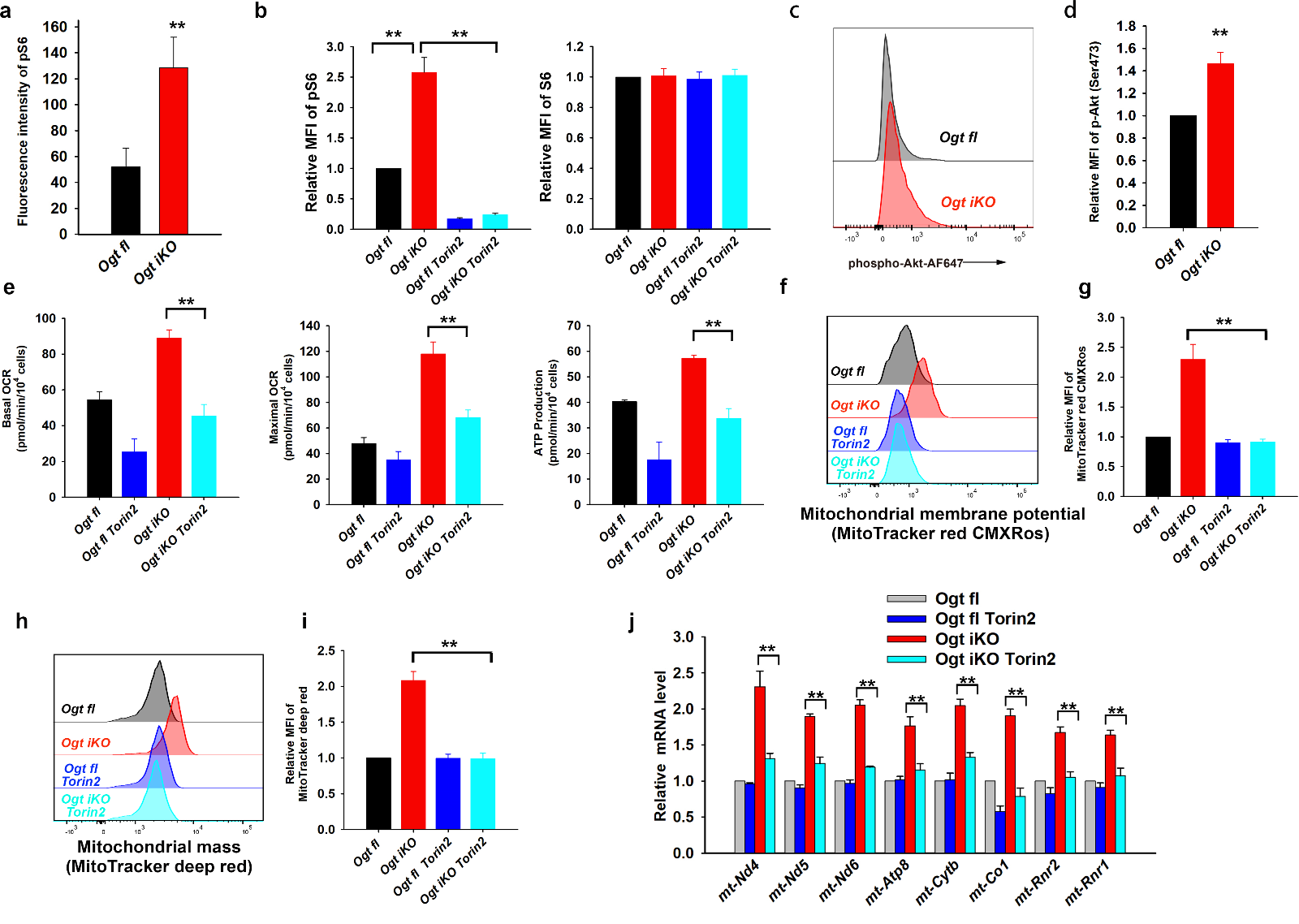

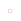

**Extended Data Fig. 7 | mTOR inhibition rescues mitochondrial dysfunction of OGT-deficient mESCs.**

**a,** Fluorescent intensity of phosphorylation of ribosomal protein S6 in *Ogt fl* and *Ogt iKO* mESCs, shown in Fig. 4b.

**b,** Quantification of the data shown in Fig. 4c, showing relative MFI (mean fluorescent intensity) of phospho-S6 and total S6 ribosomal protein, assessed by flow cytometry, in *Ogt fl* and *Ogt iKO* mESCs.

**c,** Flow cytometry analysis of phospho-Akt (Ser473) in *Ogt fl* and *Ogt iKO* mESCs treated without or with 4-OHT respectively for 6 days. Data are representative of three biological replicates.

**d,** Quantification of the data shown in **c**, showing relative MFI (mean fluorescent intensity) of phospho-Akt (Ser473), assessed by flow cytometry, in *Ogt fl* and *Ogt iKO* mESCs.

**e,** The basal OCR, maximal OCR and ATP production in *Ogt fl* and *Ogt iKO* mESCs treated with or without Torin2 for 6 days. Data are shown as mean ± SD (N=3).

**f,** Torin2 inhibits the increase in mitochondrial membrane potential in *Ogt iKO* mESCs. Representative histogram of MitoTracker red CMXRos fluorescence in *Ogt fl* and *Ogt iKO* mESCs treated with or without Torin2 (25nM) for 6 days.

**g,** Relative MFI (mean fluorescent intensity) of MitoTracker red CMXRos shown in **d**. Data are shown as mean ± SD (N=3).

**h,** Torin 2 inhibits the increase in mitochondrial mass in *Ogt iKO* mESC. Representative histogram of MitoTracker deep red fluorescence in *Ogt fl* and *Ogt iKO* mESCs treated with or without Torin2 (25nM) for 6 days.

**i,** Relative MFI (mean fluorescent intensity) of MitoTracker deep red shown in **f**. Data are shown as mean ± SD (N=3).

**j,** qRT-PCR analysis of mitochondrial DNA-encoded gene transcript levels in *Ogt fl* and *Ogt iKO* mESCs treated with or without Torin2 for 6 days. The expression level of mRNA is shown relative to the level in control mESCs. Data are shown as mean ± SD (N=3).

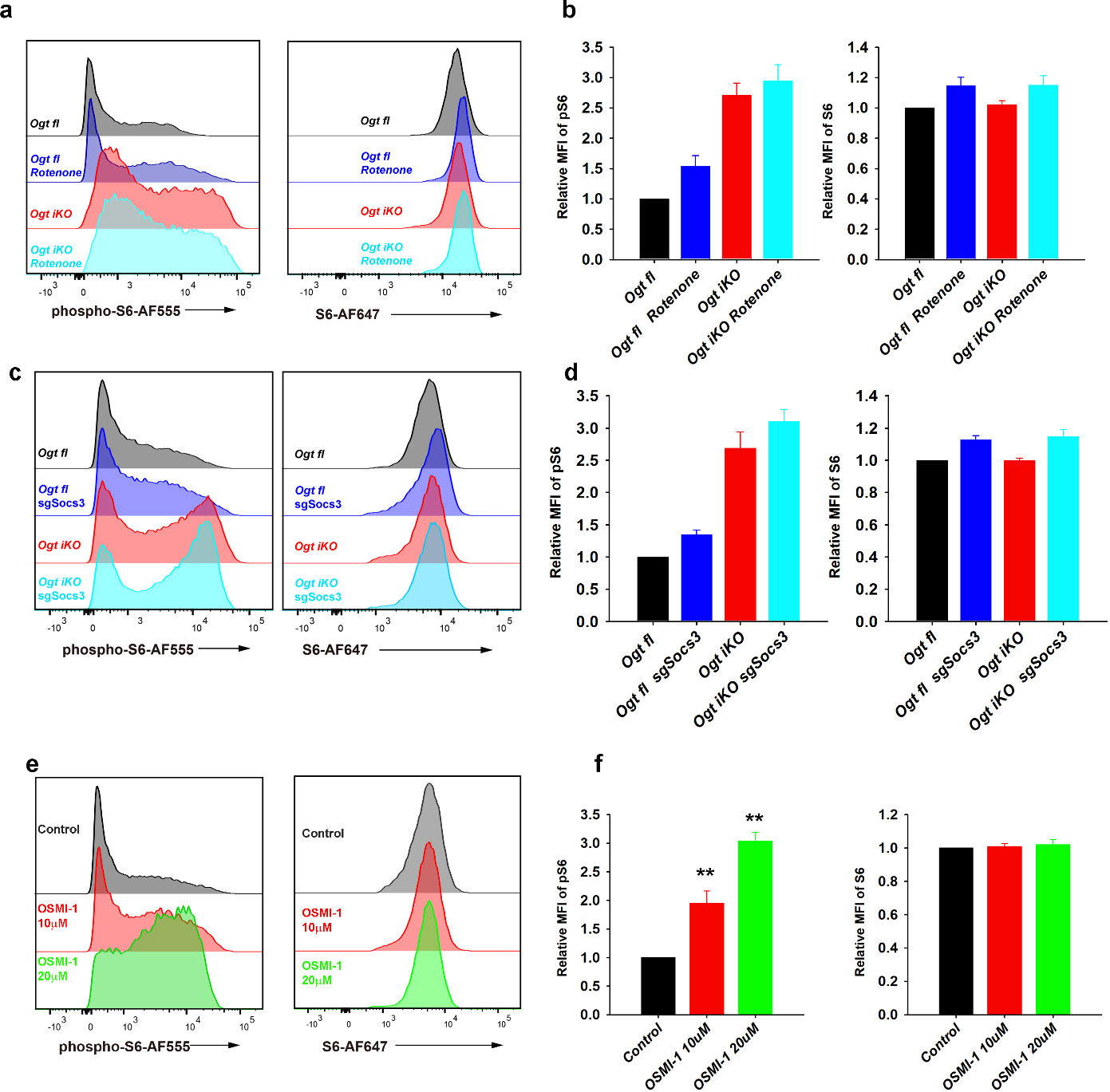
**Extended Data Fig. 8 | mTOR is hyperactivated in OGT-deficient mESCs.**

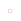

**a,** Flow cytometry analysis of phospho-S6 (*left panel*) and total S6 (*right panel*) ribosomal protein in *Ogt fl* and *Ogt iKO* mESCs treated with or without Rotenone (75 nM) for 6 days. Data are representative of three biological replicates.

**b,** Relative MFI (mean fluorescent intensity) of phospho-S6 and total S6 ribosomal protein shown in **a**. Data are shown as mean ± SD (N=3).

**c,** Flow cytometry analysis of phospho-S6 (*left panel*) and total S6 (*right panel*) ribosomal protein in *Ogt fl* and *Ogt iKO* mESCs expressing sgRNAs targeting *Socs3* for 6 days. Data are representative of three biological replicates.

**d,** Relative MFI (mean fluorescent intensity) of phospho-S6 and total S6 ribosomal protein shown in **c**. Data are shown as mean ± SD (N=3).

**e.**Flow cytometry analysis of phospho-S6 (*left panel*) and total S6 (*right panel*) ribosomal protein in *Ogt fl* mESCs treated without or with the OGT inhibitor OSMI-1 respectively for 3 days. Data are representative of three biological replicates.

**f.** Relative MFI (mean fluorescent intensity) of phospho-S6 and total S6 ribosomal protein shown in **e**. Data are shown as mean ± SD (N=3).

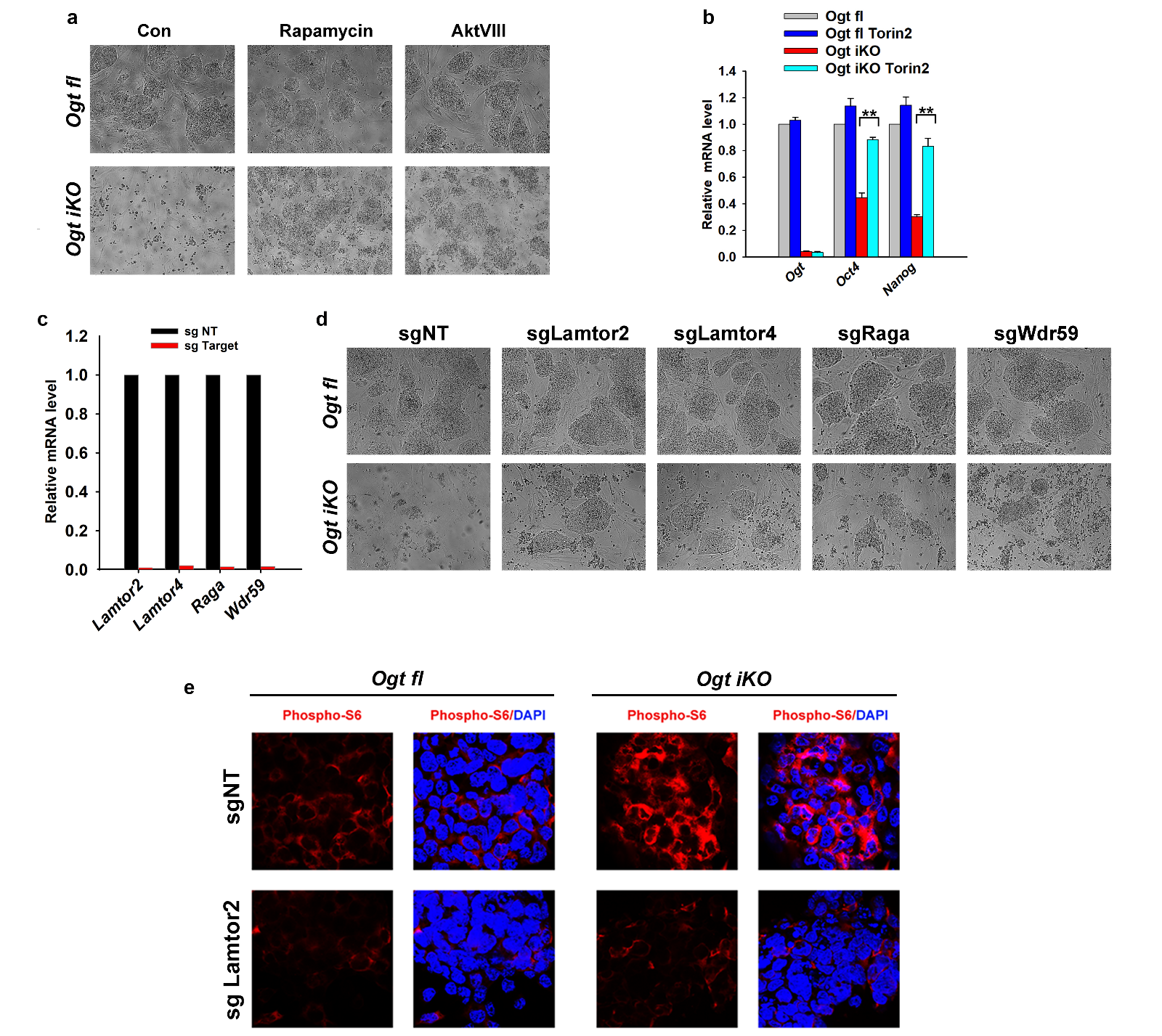

**Extended Data Fig. 9 | mTOR inhibition enabled survival of OGT-deficient mESCs.**

**a,** Phase contrast images of *Ogt fl* and *Ogt iKO* mESCs treated with or without 4-OHT and rapamycin or AktVIII for 8 days.

**b,** qRT-PCR analysis of *Ogt*, *Oct4 and Nanog* transcript levels in *Ogt fl* and *Ogt iKO* mESCs treated with or without Torin2 for 6 days. The expression level of mRNA is shown relative to the level in control mESCs. Data are shown as mean ± SD (N=3).

**c,** Individual sgRNAs efficiently deplete the indicated gene products in *Ogt fl* mESCs after puromycin selection for 7 days.

**d,** Phase contrast images of *Ogt fl* and *Ogt iKO* mESCs expressing non-targeting sgRNA (sgNT) or sgRNAs against the indicated genes and treated with or without 4-OHT for 8 days.

**e,** Immunohistochemistry of *Ogt fl* and *Ogt iKO* mESCs expressing non-targeting sgRNA (sgNT) or sgRNAs against Lamtor2 and treated with or without 4-OHT for 6 days. Cells were stained with antibody against phospho-S6 ribosomal protein. Nucleus staining: DAPI (blue). Scale bar: 10 μm.

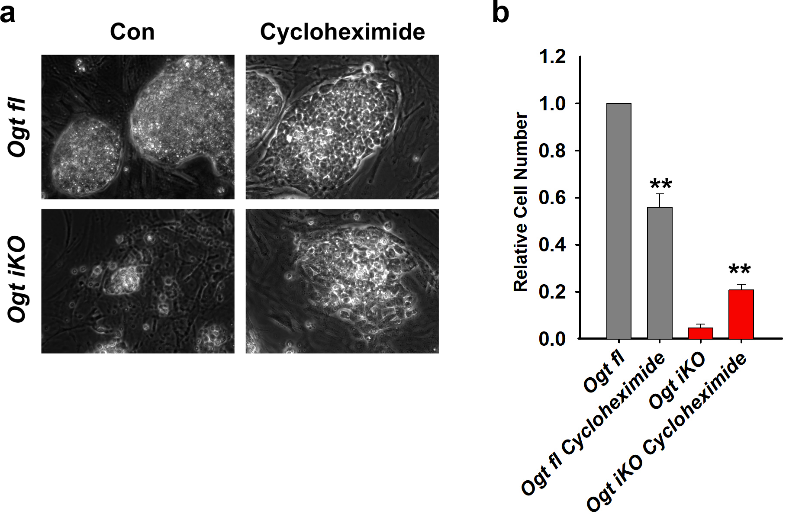

**Extended Data Fig. 10 | Protein synthesis inhibitor cycloheximide rescued survival of OGT-deficient mESCs.**

**a,** Phase contrast images of *Ogt fl* and *Ogt iKO* mESCs treated with or without 4-OHT and cycloheximide 50nM for 8 days.

**b,** Relative cell numbers of *Ogt fl* and *Ogt iKO* mESCs treated with or without 4-OHT and cycloheximide 50nM for 8 days.

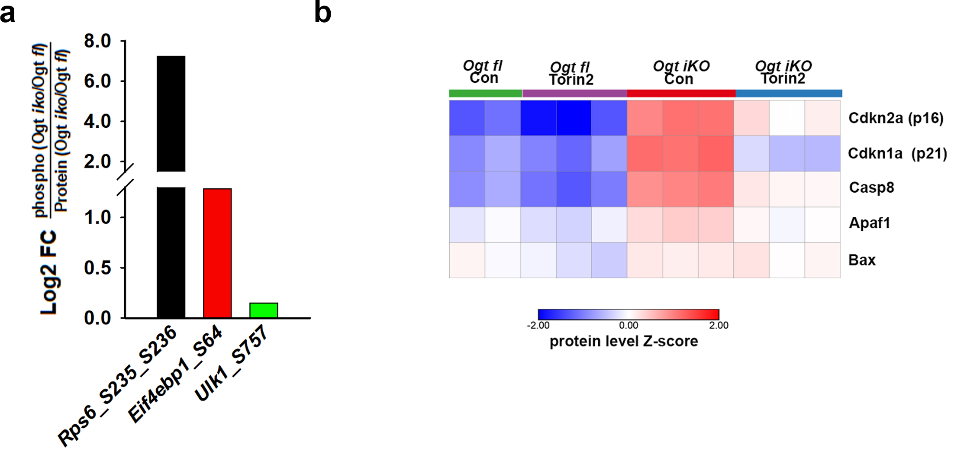

**Extended Data Fig. 11 | Phosphoproteomic analysis**

**a,** Changes in phosphopeptide and protein levels of mTOR activity markers in *Ogt fl* and *Ogt iKO* mESCs as determined by quantitative proteomics.

**b,** Changes in protein levels of *Casp8*, *Bax*, *Apa1* in *Ogt fl* and *Ogt iKO* mESCs as determined by quantitative proteomics.

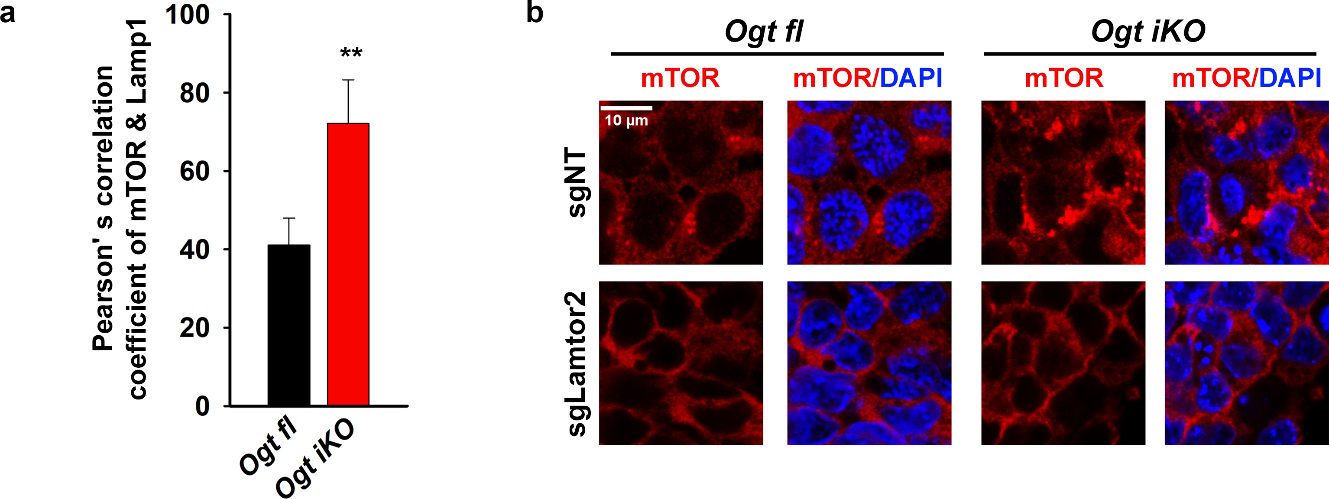

**Extended Data Fig. 12 | OGT deficiency promotes lysosomal translocation of mTOR by increasing proteasome activity.**

**a,** Statistical analysis of colocalization of mTOR and LAMP1 shown in Fig. 6a by Pearson's correlation coefficients. Images of 100 cells were quantified and data are shown as mean ± SD.

**b,** Immunohistochemistry of *Ogt fl* and *Ogt iKO* mESC expressing non-targeting sgRNA (sgNT) or sgRNAs against Lamtor2. Cells were stained with antibody against mTOR. Nucleus staining: DAPI (blue). Scale bar: 10 μm.

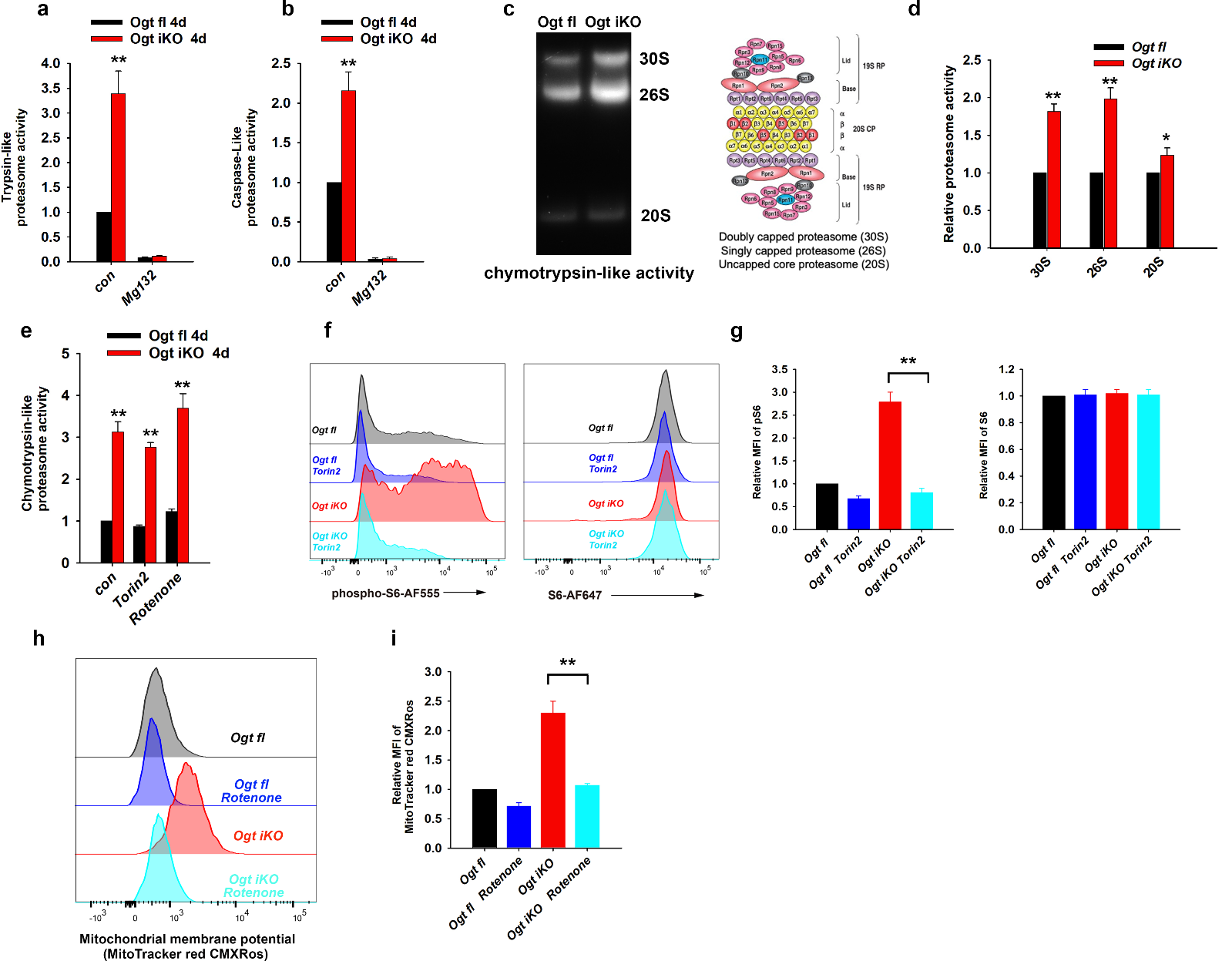

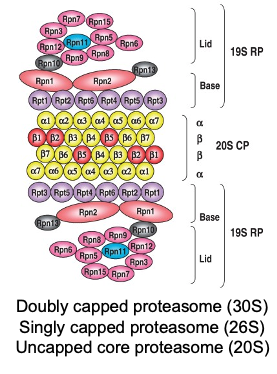

**Extended Data Fig. 13 | Proteasome activity is enhanced in OGT-deficient mESCs.**

**a-b,** Trypsin-like (**a**) and Caspase-like (**b**) proteasome activity in *Ogt fl* and *Ogt iKO* mESCs. Data are shown as mean ± SD (N=3).

**c,** Native gel analysis of chymotrypsin-like proteasome activity in *Ogt fl* and *Ogt iKO* mESCs.

**d,** Quantification of chymotrypsin-like proteasome activity shown in **c**. Data are shown as mean ± SD (N=3).

**e,** Chymotrypsin-like proteasome activity in *Ogt fl* and *Ogt iKO* mESCs treated with or without Torin2 (25 nM) or Rotenone (75 nM). Data are shown as mean ± SD (N=3).

**f,** Flow cytometry analysis of phospho-S6 (*left panel*) and total S6 (*right panel*) ribosomal protein in *Ogt fl* and *Ogt iKO* mESCs treated with or without Torin2 (25 nM). Data are representative of three biological replicates.

**g,** Relative MFI (mean fluorescent intensity) of phospho-S6 and total S6 ribosomal protein shown in **c**. Data are shown as mean ± SD (N=3).

**h,** Representative histogram of MitoTracker red CMXRos fluorescence in *Ogt fl* and *Ogt iKO* mESCs treated with or without Rotenone (75 nM). Data are representative of three biological replicates.

**i,** Relative MFI (mean fluorescent intensity) of MitoTracker red CMXRos shown in **h**. Data are shown as mean ± SD (N=3).

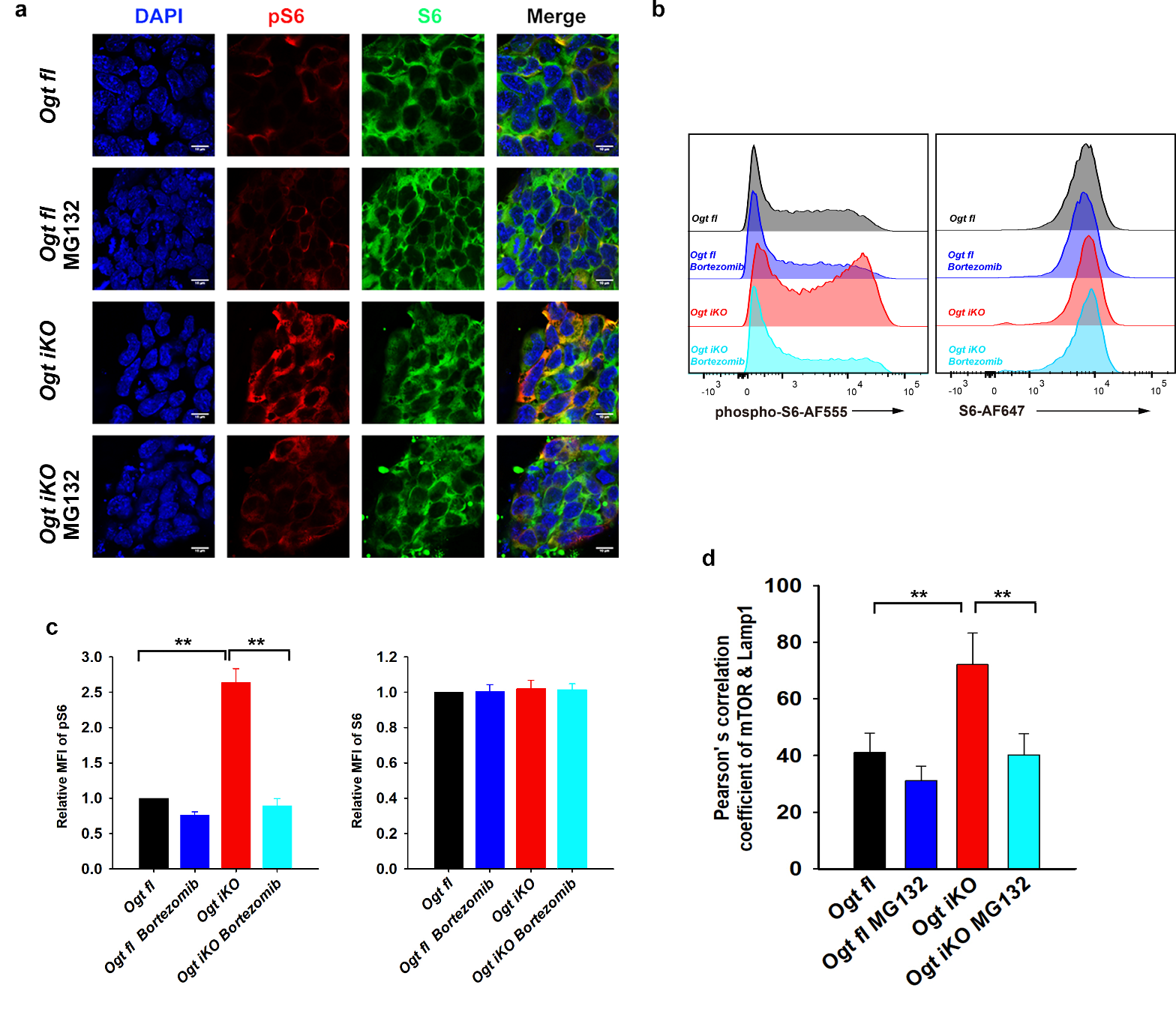

**Extended Data Fig. 14 | OGT deficiency promotes mTOR activation by increasing proteasome activity.**

**a,** Immunohistochemistry of *Ogt fl* and *Ogt iKO* mESCs treated without or with 4-OHT respectively for 6 days and then with or without 10 μM MG132 for 2.5 hours. Cells were stained with antibody against phospho-S6 and S6 ribosomal protein. Nucleus staining: DAPI (blue). Scale bar: 10 μm.

**b,** Flow cytometry analysis of phospho-S6 (left panel) and total S6 (right panel) ribosomal protein in *Ogt* *fl* and *Ogt* *iKO* mESCs treated without or with 4-OHT respectively for 6 days and then with or without 1 μM Bortezomib for 2.5 hours.

**c,** Relative MFI (mean fluorescent intensity) of phospho-S6 and total S6 ribosomal protein shown in **b**. Data are shown as mean ± SD (N=3).

**d,** Statistical analysis of colocalization of mTOR and LAMP1 shown in Fig. 6f by Pearson's correlation coefficients. Images of 100 cells were quantified and data are shown as mean ± SD.

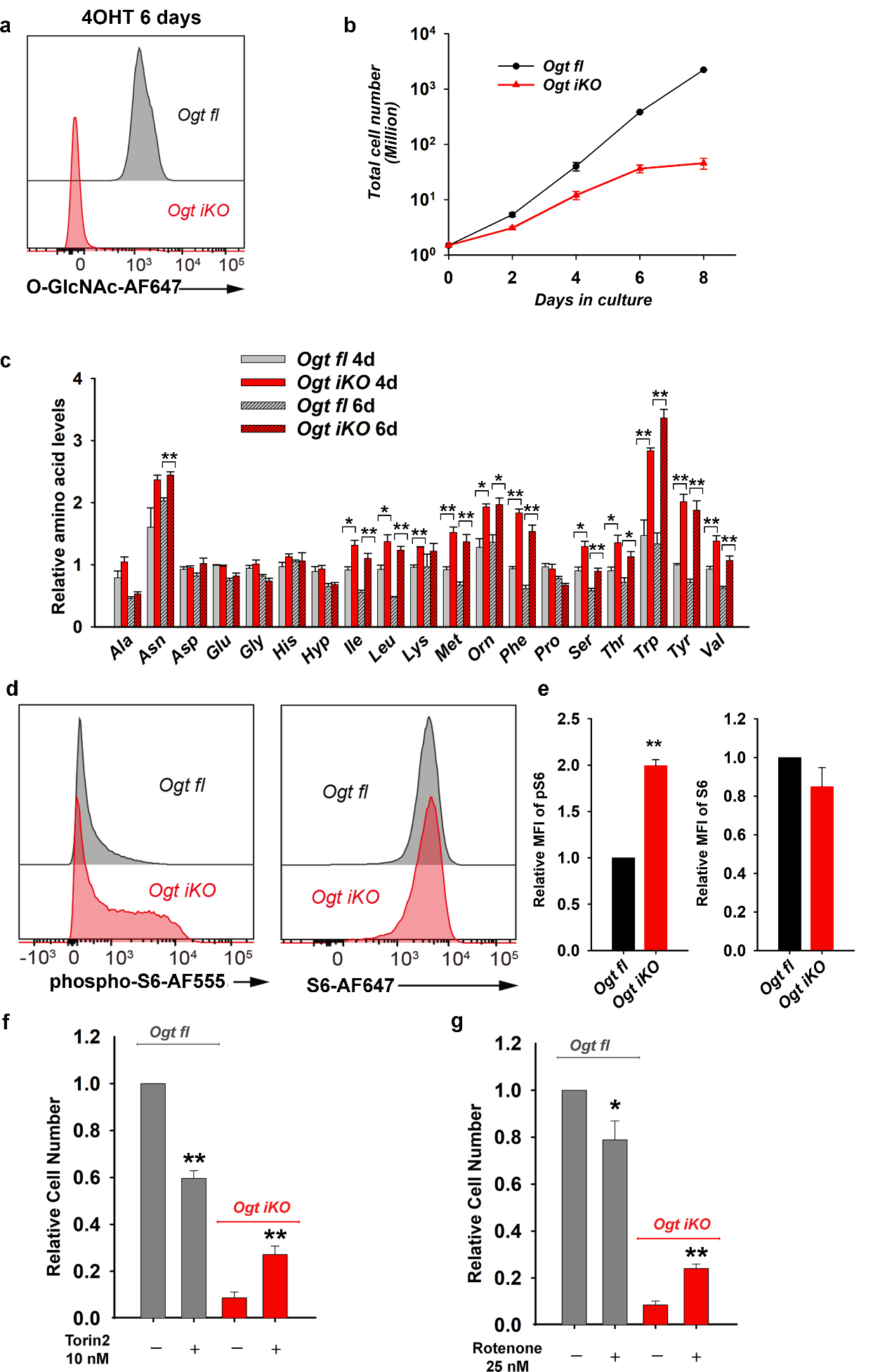

**Extended Data Fig. 15** | ***Ogt*-deficient T cells show increased amino acid levels and increased mTOR activation.**

**a,** Flow cytometry analysis of *O*-GlcNAc levels in *Ogt fl* and *Ogt iKO* CD8^+^ T cells treated without or with 4-OHT respectively for 6 days. Data are representative of three biological replicates.

**b,** Cumulative growth curves of *Ogt fl* CD8^+^ T cells treated with or without 4-OHT and counted at each passage (every 2 days) until day 8. Data are shown as mean ± SD (N=3).

**c,** Relative amino acid levels in *Ogt fl* and *Ogt iKO* CD8^+^ T cells treated without or with 4-OHT respectively for 4 or 6 days. Data are shown as mean ± SD (N=3).

**d,** Flow cytometry analysis of phospho-S6 (*left panel*) and total S6 (*right panel*) ribosomal protein in *Ogt fl* and *Ogt iKO* CD8^+^ T cells treated without or with 4-OHT respectively for 6 days. Data are representative of three biological replicates.

**e,** Relative MFI (mean fluorescent intensity) of phospho-S6 and total S6 ribosomal protein shown in **d**.

**f,** Relative cell numbers of *Ogt fl* and *Ogt iKO* CD8^+^ T cells treated without or with 4-OHT respectively and with or without Torin2 (10 nM) for 8 days. Data are shown as mean ± SD (N=3).

**g,** Relative cell numbers of *Ogt fl* and *Ogt iKO* CD8^+^ T cells treated without or with 4-OHT respectively and with or without Rotenone (25 nM) for 8 days. Data are shown as mean ± SD (N=3).

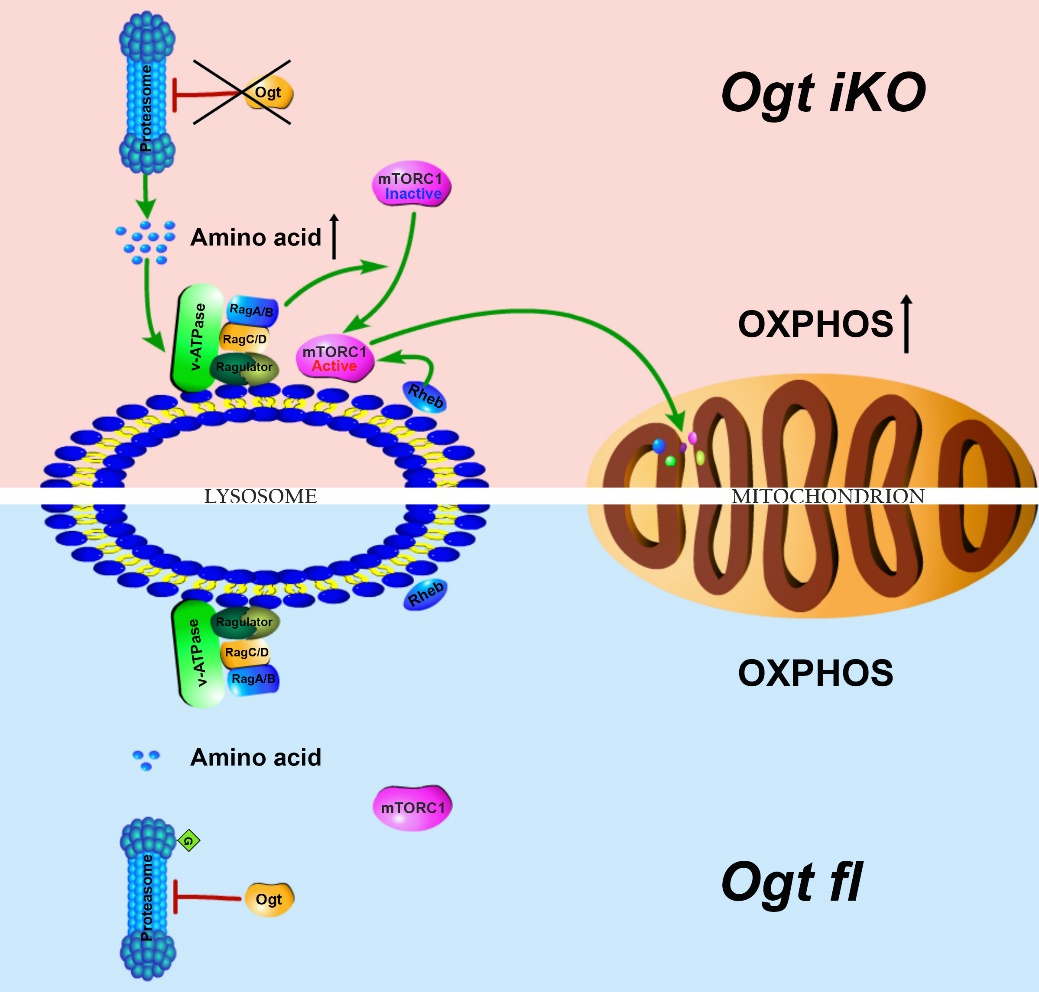
**(i) loss of OGT activity 🡪 (ii) increased proteasome activity 🡪 (iii) increased intracellular amino acid levels 🡪 (iv) hyperactivation of mTOR 🡪 (v) increased mitochondrial OXPHOS 🡪 (vi) arrested cell proliferation, eventual apoptotic cell death.**

**Extended Data Fig. 16** | **Model summarizing the findings of this study.** *Bottom*, OGT regulates cell viability by inhibiting the proteasome, thereby maintaining low levels of amino acids, mTOR activity and mitochondrial oxidative phosphorylation (OXPHOS). *Top*, in the absence of OGT, proteasome activity increases, and the consequent uncompensated increase in steady-state intracellular amino acid levels results in hyperactivation of mTOR, increased mitochondrial OXPHOS that eventually leads to decreased cell proliferation and viability.
